# Influence of historical and human factors on genetic structure and diversity patterns in peripheral populations: implications for the conservation of Moroccan trout

**DOI:** 10.1101/2020.04.06.027219

**Authors:** S Perea, M Al Amouri, EG Gonzalez, L Alcaraz, A Yahyaoui, I Doadrio

## Abstract

1. The brown trout *s.l.* has been the focus of numerous phylogeographic and conservation studies due to its socioeconomic importance, its marked genetic and phenotypic differentiation and its broad distribution range. Especially interesting evolutionary patterns are observed for populations occupying peripheral areas of a distribution range, such as in the case of the highly isolated trout populations in Morocco.
2. Continuous stocking programs may conceal natural genetic patterns, making it challenging to discern evolutionary patterns. In Morocco, trout stocking programs have been implemented to increase the genetic diversity of native populations by pooling fish of different origins in the Ras el Ma hatchery (Azrou region) and then stocking them in the different basins. In this study, phylogenetic and phylogeographic patterns, as well as genetic structure and diversity, of Moroccan trout populations were analyzed to evaluate the impact of continuous fish stocking on evolutionary processes in order to better distinguish between natural and human-mediated patterns.
3. Two mitochondrial and nine microsatellite markers were analyzed for all populations along the entire distribution range of brown trout in Morocco. Phylogenetic and phylogeographic analyses rendered two highly divergent evolutionary lineages, one comprising populations in the Drâa Basin and a second grouping the remaining Moroccan populations. Divergence of the Drâa lineage occurred during the Upper Pliocene, whilst differentiation within the second lineage coincided with the onset of the Pleistocene.
4. Genetic structuring among populations was evident. Nevertheless, populations exhibiting higher levels of genetic diversity were those affected by human-mediated processes, making it difficult to associate this diversity with natural processes. In fact, highly geographically isolated, not stocked populations showed the lowest values of genetic diversity. Although stocking management may increase the genetic diversity of these populations, it could also lead to the loss of local adaptive genotypes. Hence, current trout conservation programs should be revised.

## 1. Introduction

Biodiversity loss in freshwater systems is occurring at a faster rate than in terrestrial environments (Dudgeon et al., 2006; Strayer & Dudgeon, 2010). Exploitation of water resources, habitat alterations, mainly due to water extraction for agriculture, hydraulic infrastructure construction and urban water use, along with the introduction of exotic species, is exacerbating this loss of freshwater biodiversity (Dudgeon et al., 2006). In the Mediterranean region, climate models predict an increase in temperature and a decrease in precipitation that will decrease the availability of water resources and intensify the effects of increasing human pressure upon freshwater ecosystems (García-Ruiz, López-Moreno, Vicente Serraron, Lasanta-Martínez & Beguería, 2011; Guiot & Cramer, 2016). Populations located in the periphery of the species distribution ranges would be especially vulnerable to these global warming effects, especially in the southern periphery, that could increase the extinction risk of these populations (Gibson, van der Marel & Starzomski, 2009). Within this context, freshwater fish faunas are considered useful indicators of trends in aquatic ecosystems due to their intrinsic characteristics such as being at the top of food webs or their long longevity and high mobility (Li, Zheng & Liu, 2010; Estevez et al., 2017). Some groups are especially good indicators of water quality due to their specific ecological and habitat requirements. For example, salmonids (e.g. trout, salmon and char) have very restricted ecological requirements such as cold, clean and well-oxygenated waters and, therefore, are highly sensitive to habitat change (Almodovar, Nicola, Ayllon & Elvira, 2012; Merrian, Fernandez, Petty & Zegre, 2017; Young et al., 2018).

Within salmonids, the brown trout is distributed widely across Europe, North Africa and western Asia. Morocco constitutes the southwestern limit of the brown trout distribution range and suitable habitats for these organisms in this region are found at higher altitudes than in the northern latitudes. Consequently, these southern peripheral Moroccan trout populations have a fragmented distribution and are mainly located in headwaters of high-altitude rivers, in most cases only in the sources of these rivers, and in oligotrophic lakes in the Atlas Mountains. Some populations are also found at lower altitudes in the Mediterranean slope of the Rifian region but only in areas characterized by steep slopes with fast currents and oxygenated waters (Pellegrin, 1924). Peripheral populations of widely distributed species, as Moroccan trout, are expected not only to be more geographically isolated, showing a strong population structure but also to have smaller effective population sizes and lower genetic diversity (see Eckert, Samis & Lougheed, 2008 for a review of genetic diversity in peripheral populations). For these reasons, knowing the genetic structure and diversity of Moroccan trout populations is vital in order to evaluate the conservation status of these populations and to design appropriate conservation strategies according to the needs and characteristics of organisms from the peripheral areas of distribution ranges; Siler, Oaks, Cobb, Ota & Brown, 2014; Thorton et al., 2017).

For these conservation strategies, parameters such as genetic structure and diversity of populations, or demographic trends, are critical to design accurate plans. The patterns of genetic structure and diversity of populations are influenced by both historical and human-mediated contemporary factors that determine changes in population size and gene flow (Vucetich & Waite, 2003; Muhfeld et al., 2017), and that, ultimately, have driven the current genetic patterns found in Moroccan trout. Therefore, to understand how the genetic structure, diversity and demography of Moroccan trout populations have been shaped along its evolutionary history, it is essential to consider on the one side the complex geomorphological landscape of the High Atlas Mountains with its rivers flowing between steep canyons and the climatic events that cause drastic floods in Morocco, that is a result of high tectonic activity occurred during the Neogene and Quaternary eras (Michard, Frizon de Lamotte, Saddiqi & Chalouan, 2008; Babault, Van den Driessche & Texeill, 2012; El Fels, Alaa, Bachnou & Rachidi, 2018), and on the other side contemporary factors related to human activity in this country. This complex geological history of Morocco has generated its highly diverse mountains and river basins, and their highly isolated streams. Therefore, a high level of genetic structure is expected in Moroccan trout since the formation of these freshwater ecosystems.

Contemporary factors related to human activity are increasingly crucial to understand current genetic structure and diversity of Moroccan trout populations. Aquatic resources are being damaged worldwide due to increased infrastructure construction and overexploitation of these resources for agricultural purposes and tourism (Tekken & Kropp, 2015; Molle & Tanouti, 2017). The effects of these activities have been especially detrimental in Morocco, where the annual total renewable water resources per capita is six-fold lower than the global average, and water shortages are frequent (Doukkali, 2005; Tekken & Kropp, 2015). Overfishing of trout populations, a direct consequence of Morocco’s increased infrastructure and tourism, is another contemporary factor affecting the genetic structure and diversity of populations (Almodovar & Nicola, 2004; Sánchez-Hernández, Shaw, Cobo & Allen, 2016). The genetic erosion of populations due to the introduction of non-native lineages, a common worldwide practice for sport fishing, is another threat for trout populations in general (Almodovar, Nicola, Elvira & García-Marín, 2006; Vera, García-Marín, Martínez, Araguas & Bouza, 2013; Arthington, Dulvy, Gadstone & Winfield, 2016). Introgression of different genetic lineages due to stocking may conceal natural genetic patterns as a consequence of population structure erosion and the introduction of non-native populations (Petereit et al., 2018; Vera, Martínez & Bouza, 2018). Indeed, the brown trout is one of the most important riverine fishes, particularly for its role in local and sport fishing, which have favored massive artificial translocations of individuals among basins. Consequently, over the last several decades, populations have been mixing over entire brown trout distribution range, increasing the complexity of the evolutionary processes affecting this group (Horreo, Abad, Dopico, Oberlin & García-Vázquez, 2015; Sanz, 2017; Závorka et al., 2017).

Given their fragmented and peripheral nature, some Moroccan trout populations have drastically decreased in the last decades (Clavero et al., 2017). Since 1957, the Moroccan government has maintained a trout pool comprised of different native populations in the Ras el Ma hatchery (Azrou region). Nevertheless, little information exists about the origin and contribution of each population to the Ras el Ma hatchery stock; therefore, the impact of stocking processes on the genetic diversity and structure of wild trout populations in Morocco is unknown. The majority of studies evaluating the impact of trout stocking in southern Europe from central European populations are based on the analysis of a variant of the *lactate dehydrogenase C* (*LDH-C*) locus that was not initially present in the southern European populations (McMeel, Hoey & Ferguson, 2001; Kohout, Papousek, Sediva & Slechta, 2012). However, the management policy in effect in Morocco differs from Europe in that native Moroccan trout populations are used for stocking. Therefore, an analysis of *LDH-C* is not a viable tool to assess hybridization in these populations.

The main aims of this study are to (1) determine the phylogenetic and phylogeographic structure of Moroccan trout populations (*Salmo spp*), (2) estimate their genetic diversity and demographic parameters and (3) assess the impact of fish stocking from the Ras el Ma hatchery on wild trout populations through the analysis of their genetic variation. For this purpose, trout populations from different locations across the highlands of Morocco were studied genetically through the sequence analyses of the mitochondrial *MT-CYB* gene and D-loop region and the genotyping of nine microsatellite loci. The identification of the historical and contemporary factors driving the genetic and demographic patterns found in Moroccan trout populations will provide insight on the evolutionary processes that have occurred, or are occurring, at the southwestern limit of the distribution range of brown trout. The knowledge gained from this study may help inform relevant conservation strategies for this freshwater fish group of great socioeconomic importance.

## 2. Material and methods

### 2.1. Sampling

A total of 475 individuals of the genus *Salmo* were collected between 2009 and 2015 from 19 localities across the Mediterranean, the Atlantic and desert river basins located at the southern end of the Atlas Mountains (Figure 1, Table 1). Brown trout populations from the different basins, and samples representing all recognized lineages except for the Tigris (Bernatchez, 2001; Suárez, Bautista, Almodovar & Machordom, 2001; Bardakci et al., 2006), were included in the phylogenetic and phylogeographic analyses (Table 1). Moreover, sequences from a population in Sicily (Italy), which has been shown to be closely related to Moroccan trout populations (Tougard et al., 2018), and from three specimens belonging to the species *Salmo macrostigma* obtained from GenBank (LT617630–LT617632, including the holotype housed at the Natural History Museum of Paris; Tougard et al., 2018) were included in the phylogenetic analyses. Specimens were captured by electrofishing or mesh nets with the permission of local authorities, fin clipped and then returned to the stream, except for two or three individuals per locality, which were kept as reference specimens. Fin clips were preserved in 95% ethanol. All vouchers samples are stored at the Museo Nacional de Ciencias Naturales, CSIC, Madrid, Spain. Locations that are supposed to remain not stocked were selected based on information provided by the fishing committee of Haut Commissariat aux Eaux et Forêts et à la Lutte Contre la Désertification (HCEFLCD).

**Table 1.** Populations and sampling site locations, including the names of the corresponding river and lake basins from which samples were collected. The number (N) of individual *Salmo trutta* samples included in the mitochondrial (*MT-CYB* and D-loop) and nuclear microsatellite DNA analyses are also indicated. GeneBank accesion numbers are provided for the *MT-CYB* and D-loop sequences. (Note: awaiting for Genbank accession numbers)

| POPULATION<br>(NUMBER IN MAP) | LOCALITY | N (*MT-CYB* / *D-LOOP* / *STR*) | *MT-CYB*<br>*ACCESSION*<br>*NUMBERS* | *D-LOOP*<br>*ACCESSION*<br>*NUMBERS* |
| --- | --- | --- | --- | --- |
| **Moroccan populations** |  |  |  |  |
| **FARDA (1)** | Laou Basin /<br>Mediterranean slope | 10 / 10 / 15 |  |  |
| **KANNAR (2)** | Kannar Basin /<br>Mediterranean slope | 12 / 13 / 16 |  |  |
| **MOULOUYA (4)** | Moulouya Basin /<br>Mediterranean slope | 20 / 20 / 20 |  |  |
| **SIDI HAMZA (5)** | Ziz Basin / flowing<br>to Sahara | 40 / 23 / 50 |  |  |
| **SIDI RACHID (17)** | Sidi Rachid R.<br>Sebou Basin/<br>Atlantic Slope | 34 / 34 / 32 |  |  |
| **MIAAMI (3)** | Oum er Rbia Basin/<br>Atlantic slope | 16 / 15 / 14 |  |  |
| **MELLOUL (7)** | Oum er Rbia Basin/<br>Atlantic slope | 30 / 32 / 29 |  |  |
| **LAKHDAR (10)** | Oum er Rbia Basin/<br>Atlantic slope | 24 / 26 / 26 |  |  |
| **TESSAOUT (11)** | Oum er Rbia Basin /<br>Atlantic slope | 20 / 20 / 15 |  |  |
| **AMENGOUSS (18)** | Oum er Rbia Basin /<br>Atlantic slope | 4 / 4 / 4 |  |  |
| **AIT NACER (19)** | Oum er Rbia Basin /<br>Atlantic slope | 3 / 3 / 0 |  |  |
| **OURIKA (13)** | Tensift Basin /<br>Atlantic slope | 28 / 26 / 28 |  |  |
| **RHERAYA (14)** | Tensift Basin /<br>Atlantic slope | 14 / 14 / 13 |  |  |
| **TIFNOUTE (16)** | Souss Basin/<br>Atlantic slope | 26 / 29 / 29 |  |  |
| **M'GOUN (9)** | Drâa Basin /<br>Atlantic slope | 27 / 29 / 29 |  |  |
| **DADES (8)** | Drâa Basin /<br>Atlantic slope | 35 / 34 / 34 |  |  |
| **ISLI (6)** | Endorheic lake | 24 / 23 / 26 |  |  |
| **IFNI (15)** | Endorheic lake | 17 / 21 / 23 |  |  |
| **TAMDA (12)** | Endorheic lake | 40 / 43 / 35 |  |  |
| **Other populations** |  |  |  |  |
| **GARONA** | Garona Basin /<br>Atlantic slope<br>(Iberian Peninsula) | 6 / 9 / 0 |  |  |
| **MANDEO** | Mandeo Basin /<br>Cantabrian slope<br>(Iberian Peninsula) | 4 / 4 / 0 |  |  |
| **ARANEA** | Aranea Basin /<br>Cantabrian slope<br>(Iberian Peninsula) | 2 / 2 / 0 |  |  |
| **DUERO** | Duero Basin /<br>Atlantic slope<br>(Iberian Peninsula) | 6 / 9 / 0 |  |  |
| **TURIA** | Turia Basin /<br>Mediterranean slope<br>(Iberian Peninsula) | 5 / 9 / 0 |  |  |
| **GUADALQUIVIR** | Guadalquivir Basin /<br>Atlantic slope<br>(Iberian Peninsula) | 21 / 25 / 0 |  |  |
| **SICILY** | Sicily | 3 / 3 / 0 | LT617555-LT617557 | LT617612-LT617614 |
| **ADRIATIC**<br>**LINEAGE** |  | 3 / 3 / 0 | LT617523,<br>LT617528-LT617529 | LT617590,<br>LT617595-LT617596 |
| **MARMORATUS**<br>**LINEAGE (*Salmo***<br>***marmoratus*)** |  | 2 / 2 / 0 | LT617574-LT617575 | LT617616-LT617617 |
| **MEDITERRANEAN**<br>**LINEAGE** |  | 2 / 2 / 0 | LT617581-LT617582 | LT617623-LT617624 |
| **DANUBIAN**<br>**LINEAGE** |  | 3 / 3 / 0 | LT617546-LT617548 | LT617608-LT617610 |
| *Salmo macrostigma* |  | 3 / 3 / 0 | LT617630-LT617632 | LT617630-LT617632 |
| *Salmo ohridanus* |  | 2 / 2 / 0 | AY926568 | AY926569 |
| *Salmo obtusirostris* |  | 2 / 2 / 0 | JX960841 | EF469833 |

**Figure 1.**
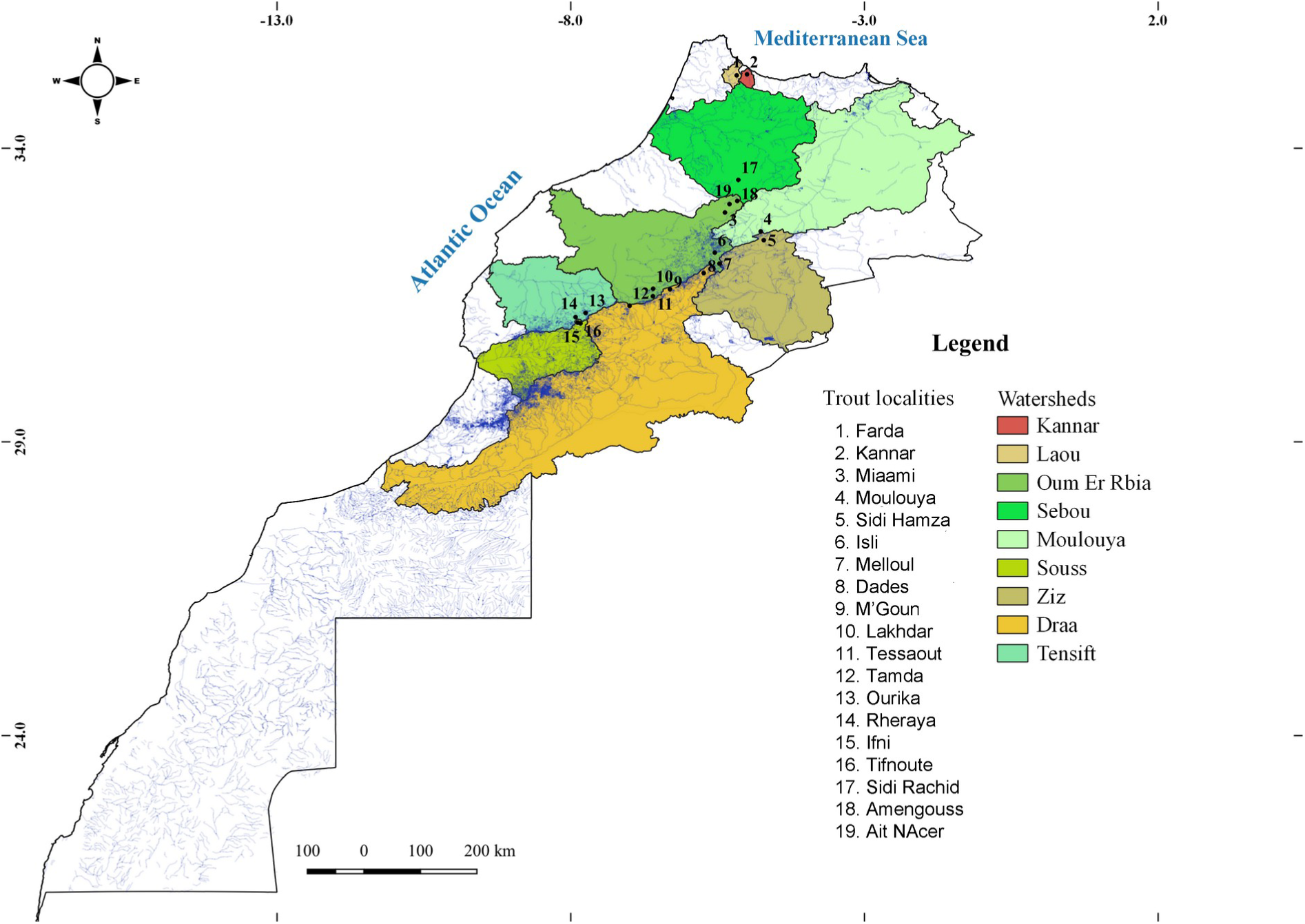
Sampling localities of *S. trutta* in Morocco.

### 2.2. Mitochondrial DNA amplification and sequencing

Total genomic DNA was extracted from fin tissue using the BioSprint15 DNA Blood Kit (Qiagen). The complete mitochondrial cytochrome *b* gene (*MT-CYB*; 1140 bp) and a fragment of D-loop, a region of the mtDNA control region (999 bp final alignment including gaps), were amplified by polymerase chain reaction (PCR). For the D-loop region, the primers LN20 (5’-ACCACTAGCACCCAAAGCTA-3’) and HN20 (5’-GTGTTAGTCTTTAGTTAAGC-3’) were used (Bernatchez & Danzman, 1993). The D-loop region could not be amplified in some samples with the above primers. Therefore, a new primer (cytb-dloop-st: 5’-ATCGGTCAAGTTGCCTCTG-3’) was designed using OLIGO 7 (Rychlik, 2007). For *MT-CYB*, the primers Glu-F and Thr-R (Zardoya & Doadrio, 1998) were used. The PCR reaction consisted of a final volume of 25μl and included 0.2μM of each primer, 0.25μM of each DNTP, 10X PCR buffer, 1.5 U Taq polymerase (5 PRIME) and 40 ng of genomic DNA. The following thermocycling conditions were used: initial denaturation at 95 °C for 5 min followed by 40 cycles of denaturation at 94 °C for 45s, annealing at 52 °C for D-loop and 48 °C for *MT-CYB* for 45s and extension at 72 °C for 90s, and a final extension at 72 °C for 10 min. Amplified DNA fragments were checked on 1.5% agarose gels, purified using ExoSAP-IT (USB) and directly sequenced on an ABI 3730XL DNA Analyzer by Macrogen Europe Inc. (http://www.Macrogen.com).

### 2.3. Microsatellite amplification and genotyping

A total of 445 samples were genotyped for nine previously described dinucleotide microsatellite markers: STR60 and STR15 (Estoup et al., 1993), STR85 and STR543 (Presa & Guyomard, 1996), STR131 (Estoup et al., 1998), STR591 and STR541 (Estoup et al., 2002), SSA103 (Thorsen et al., 2005) and SSA100 (Unpublished, see Giger et al., 2006 supplementary material). Different multiplex reactions were amplified: STR591+STR541, STR543+SSA100 and SSA103+STR60+STR131+STR15. Locus STR85 was amplified separately because it required a different annealing temperature. DNA amplifications were performed in 20 μl reactions that contained 2X QIAGEN multiplex PCR Master Mix*, following manufacturer conditions. Forward primers were labeled with fluorescent dyes, and reactions were run on a Veriti Thermal Cycler (Applied Biosystems). PCR products were checked on 2% agarose gels. The amplified fragments were sequenced with an ABI 3730 DNA sequencer (Applied Biosystems) by Secugen (Madrid, Spain), and allele sizes were assigned using the program GeneMapper v3.7 (Applied Biosystems).

### 2.4. Phylogenetic, phylogeographic and divergence time estimation analyses

*MT-CYB* and D-loop sequences were aligned in Geneious (Kearse et al., 2012). The best-fit model of evolution for each marker (*MT-CYB* and D-loop) and for each codon position of the *MT-CYB* gene was estimated via Akaike Information Criterion in PartitionFinder v.1.1.1 (Lanfear, Calcott, Ho & Guindon, 2012). The best scheme partition is represented in Table S1 (Supporting Information). The species *Salmo salar* and *Salmo ohridanus* were used as outgroups. Bayesian inference (BI) was performed in MrBayes 3.2 (Ronquist et al., 2012). For the analysis, two simultaneous runs were performed for 107 generations, each one with four MCMC chains, sampling every 100 generations. Convergence was checked with Tracer v.1.7. (Rambaut, Drummond, Xie, Baele & Suchard, 2018). After discarding the first 10% of generations as burn-in, the 50% majority rule consensus tree and the posterior probabilities were obtained. Maximum Likelihood (ML) reconstruction was conducted with RAxML in the Trex-online server using the substitution model GTRGAMMAI and the rapid bootstrapping algorithm (Stamatakis, 2006; Stamatakis, Blagojevic, Nikopoulos & Antonopoulos, 2007). We assessed node confidence using 10,000 non-parametric bootstrap replicates. Uncorrected-p genetic distances were also estimated in MEGA v.7 (Kumar et al., 2016) in order to check for differences between and within populations. To assess the phylogeographic history of the genus *Salmo* in Morocco, we reconstructed a haplotype network with *MT-CYB* and D-loop sequences independently to evaluate the shallow relationships among closely related haplotypes. The haplotype network was constructed using POPART (Leigh & Bryant, 2015), and the median-joining algorithm was used following default parameters, as recommended for multiple state data (Bandelt, Forster & Röhl, 1999).

Divergence time estimation was carried out using the concatenated mitochondrial dataset in BEAST v.1.8.3 (Drummond, Suchard, Xie & Rambaut, 2012). Only one individual per brown trout population was included in the analysis. *Salmo ohridanus* and *Salmo salar* were also included in the analyses. The phylogenetic and phylogeographic analyses performed in this study showed the artificial origin of the Tamda population; therefore, this population was excluded from this analysis. Some of the other studied populations could be affected by restocking from the Ras el Ma hatchery; however, on the basis of a native population hypothesis and on the lack of knowledge about its impact in native populations, all remaining populations were taken into account in the divergence time estimation analysis. A random local clock, which has rate heterogeneity into account, and a Birth-Death speciation model were used in the analysis. Due to the lack of old and reliable fossils, the molecular clock was calibrated using two different biogeographical points. Given that the *S. ohridanus* lineage was included in the phylogenetic analysis, the first point was the age of the formation of Lake Ohrid in the Pliocene, estimated to have occurred between 5-2 Mya (Albrecht & Wilke, 2008). The second point was the formation of Isli Lake in the Early-Middle Pleistocene (Ibouh, Michard, Charrière, Benkkadour & Rhoujjati, 2014). These calibration points were incorporated into the analysis as normal priors with a distribution encompassing the range of the age estimated for the formation of both lakes. A secondary calibration point based on the divergence of *Salmo salar* and *S. trutta* around 14-17 Mya (Horreo, 2017) was also considered in the analysis. MCMC analyses were run for 400 million generations, with parameters logged every 10,000 generations. Default settings were used for the remaining parameters. Convergence of the analysis was evaluated in Tracer v1.7 (Rambaut et al., 2018), and results were summarized in TreeAnotator v.1.8.3 (Drummond et al., 2012).

### 2.5. Genetic structure analyses of the microsatellite and mitochondrial data

Two Bayesian clustering methods were used to determine the population structure of *Salmo trutta* based on microsatellites. The number of populations (*K*) with the highest posterior probability (mean lnProb(D)) under an admixture model was calculated with the program STRUCTURE 2.3 (Pritchard, Stephens & Donelly, 2000). MCMC simulations consisted of 104 burn-in iterations followed by 106 sample iterations. Each simulation was run 10 times, exploring K-values from 1 to 17 (total number of analyzed populations). The most likely number of homogeneous clusters (best value of K) was assessed through the modal value of delta, Δ*K*, using the online application STRUCTURE HARVESTER (Evanno, Regnaut & Goudet, 2005; Earl & VonHoldt, 2012). CLUMPAK software (Kopelman, Mayzel, Jakobsson, Rosenberg & Mayrose, 2015) was used to summarize results from the 10 independent runs, and the results were represented graphically using DISTRUCT (Rosenberg, 2004). Complementary Bayesian clustering analysis was performed in TESS v.3 (Caye, Deist, Martins, Michel & François, 2016), considering both CAR and BYM admixture models, which address spatial autocorrelation and complex spatiotemporal processes (Durand, Jay, Gaggioti & François, 2009). Exploratory data analyses for different numbers of K were run to evaluate the maximum number of clusters (Kmax), setting 50,000 sweeps, a burn-in of 10,000 and starting with a neighbour joining tree. The model with the lowest deviance information criterion (DIC) value and that stabilized at the lowest number of clusters was chosen as the one that best explained the genetic variation in the data for both admixture models.

To assess the relative contribution of genetic variation to structure within and among Moroccan trout populations, we performed an analysis of molecular variance (AMOVA) based on microsatellites and mitochondrial markers, implemented in Arlequin v.3.1.5.2 (Excoffier & Licher, 2010). Genetic differentiation among populations was addressed through Φ_ST_ pairwise comparisons (mitochondrial; Hudson, Slatkin & Maddison, 1992) and through the fixation index (*F*_ST_) (microsatellites; Weir and Cockerham, 1984) using Arlequin v.3.1.5.2 (Excoffier & Licher, 2010). Statistical significance of Φ_ST_ and *F*_ST_ estimates was determined by 1,000 permutations of individuals among populations. Significant deviations from the null hypothesis of no differentiation were assessed with 10,000 permutation tests. As multiple paired tests were performed, p-values were adjusted by Bonferroni’s correction (Rice, 1989). For microsatellites, genetic relationships among samples were also examined using a factorial correspondence analysis (FCA) implemented in GENETIX v.4.05 (Belkhir, Borsa, Chikhi, Raufaste & Bonhomme, 2004), which places the individuals in a two-dimensional space according to the degree of similarity in their allelic state.

### 2.6. Mitochondrial and Microsatellite genetic diversity analysis

The following genetic parameters of diversity for *MT-CYB* and D-loop were estimated using the software DNAsp v5.0 (Librado & Rozas, 2009): number of haplotypes (h), nucleotide diversity (π), haplotype diversity (*H*_D_), number of polymorphic sites (*S*) and number of pairwise differences (*K*). For microsatellites, the presence of null alleles was estimated using the Oosterhout’s estimator as implemented in MICRO-CHECKER 2.2.3 (Oosterhout, Weetman & Hutchinson, 2006). Allelic frequencies at the different loci were estimated with the program FreeNA (Chapuis & Estoup, 2007). Microsatellite genetic diversity was quantified for each locus and population based on the average number of alleles per locus (*N*A), number of alleles standardized to those of the population sample with the smallest size (*NAR*) (Nei & Chesser, 1983) and the observed (*H*O) and expected (*H*E) heterozygosities (Nei, 1978) using GENETIX 4.05 (Belkhir et al., 2004) and FSTAT (Goudet, 2001). Deviations from Hardy-Weinberg (HW) proportions were tested using the exact probability test for multiple alleles (Guo & Thompson, 1992), available on the web-based version of GENEPOP 4.2 (Rousset, 2008), at each locus for each population and over all loci for each population. Genotypic linkage disequilibrium between loci pairs was estimated by Fisher’s exact test with the web-based version of GENEPOP 4.2 (Rousset, 2008).

### 2.7. Population size changes and gene flow based on mitochondrial and microsatellites markers

To evaluate population size changes based on mitochondrial data, deviations from a model of mutation-drift equilibrium were tested for both mitochondrial markers using the Fu’s *FS* (Fu, 1997), *R2* (Ramos-Onsins & Rozas, 2002) and Tajima’s *D* (Tajima, 1989) neutrality tests as implemented in DNAsp v5.0 (Librado & Rozas, 2009). To assess demographic trends, mismatch analyses were carried out in DNAsp v5.0 (Librado & Rozas, 2009). Initial values were set at θ_0_ = 0 and θ_1_ = 99,999. The fit of the data to the sudden demographic expansion model was tested by the probability of obtaining smaller raggedness values (*r*) than those observed under coalescent algorithm simulations over 1000 pseudo-replications and with no recombination (Rogers & Harpending, 1992). Gene flow among Moroccan trout populations based on mitochondrial markers was estimated through the virtual number of migrants (*Nm*) exchanged among populations per generation (Slatkin & Barton, 1989) using Arlequin v3.5.2.1 (Excoffier & Lischer, 2010).

Possible decreases in effective population size on the basis of microsatellites were assessed using the software BOTTLENECK (Piry, Luikart & Cornuet, 1999). Analyses were performed considering three mutation models: 1) infinitive allele (IAM), 2) stepwise mutation (SMM) and 3) two-phase (TPM) considering 70% stepwise and 30% variable mutations. A one-tailed Wilcoxon signed-rank test for statistical detection of Hardy-Weinberg heterozigosity excess was performed, as this test is more accurate in cases with a low number of polymorphic loci (Luikart, Allendorf, Cornuet & Sherwin, 1998). Estimation was based on 10,000 replicates. The mode-shift test (Luikart et al., 1998) in BOTTLENECK was also conducted to determine whether the observed distribution of allele frequencies among Moroccan trout populations differed from expectation under drift-mutation equilibrium (L-shaped distribution). As with the mitochondrial markers, gene flow among Moroccan trout populations based on microsatellites was estimated through the virtual number of migrants (*Nm*) exchanged among populations per generation (Slatkin & Barton, 1989) using Arlequin v3.5.2.1 (Excoffier & Lischer, 2010).

## 3. Results

### 3.1. Phylogenetic and phylogeographic analyses

We obtained a total of 462 individuals for the complete *MT-CYB* gene (1140bp) and 472 individuals for a fragment of the mitochondrial D-loop region (999 bp final alignment including gaps). The concatenated mtDNA dataset used for phylogenetic analyses comprised a total of 387 individuals and 2139 bp (Table 1). Bayesian and Maximum Likelihood topologies for this dataset were concordant (Figure 2). Phylogenetic analyses revealed that Moroccan trout populations are not monophyletic and the presence of three main highly supported clades. Clade I included samples from the Moroccan Drâa Basin (Dades and M’Goun tributaries, corresponding to *S. multipunctatus*), the Adriatic, Danubian and Mediterranean lineages of *S. trutta*, *S. marmoratus* and Iberian populations from the Guadalquivir Basin. Clade II comprised the other Moroccan populations analyzed along with the Algerian *S. macrostigma* (holotype: GenBank mtDNA sequence LT617631) and populations from Sicily (Figure 2). Clade II was in turn subdivided into several well-supported subclades (Figure 2). Thus, Isli Lake (corresponding to *S. viridis*) and Rifian (Farda and Kannar) populations resolved as monophyletic groups. Populations within the Oum er Rbia Basin were not monophyletic as individuals from one of its tributaries (Lakhdar R.) were included in two different subclades: one comprised of populations from other Oum er Rbia basin tributaries and the other consisting of the endorheic Sidi Hamza population (Ziz Basin) and the Mediterranean Moulouya basin population. This latter population was also recovered as polyphyletic as some Moulouya individuals were nested within the Oum er Rbia, Ourika and Ifni–Tifnoute subclades. Within the subclade comprising populations of the Oum er Rbia basin tributaries, the population Tessaout constituted an independent and well-supported clade (data not shown; pp = 1.0, bootstrap = 95). Sidi Rachid population (Sebou Basin) was clustered within the Sidi Hamza subclade. Populations from the Tensift Basin (adscribed to *S. pellegrini*) were not monophyletic, as populations from its two tributaries, Ourika and Rheraya, did not clustered together. Although Rheraya constituted a well differentiated clade, some of its individuals clustered within the Oum er Rbia basin subclade. Ourika individuals grouped with some from the Moulouya River (River is herein abbreviated as R.). The Ifni lake population (corresponding to *S. akairos*) was closely related to the population found in the Tifnoute R., a tributary of the Souss Basin that is geographically close to Ifni Lake, and some representatives of the Moulouya and Tamda populations. Brown trout from the Iberian Duero Basin constituted and independent lineage (Clade III). Phylogenetic relationships among the three clades were not fully solved.

**Figure 2.**
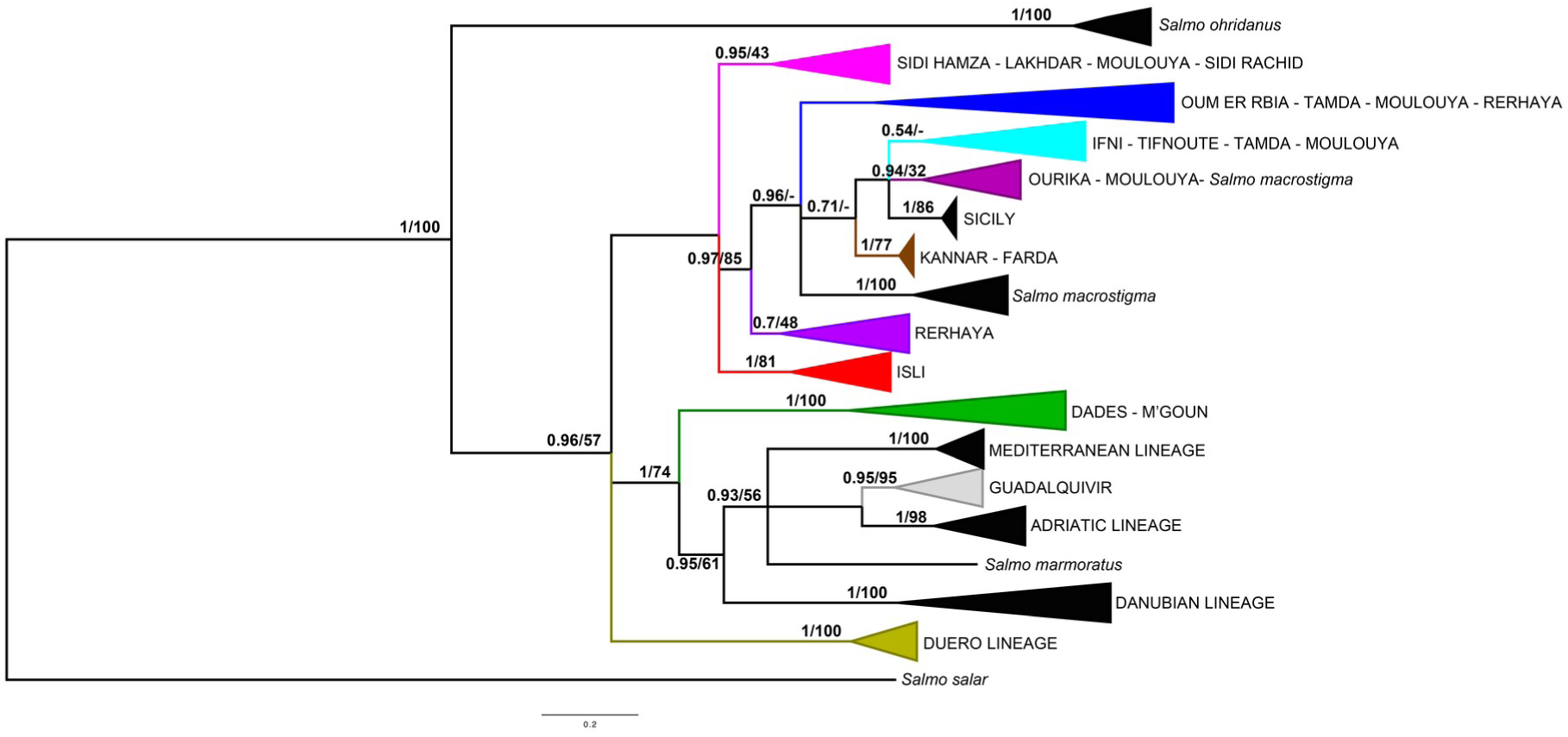
Bayesian phylogenetic tree of Moroccan trout populations based on the analysis of mtDNA (*MT-CYB* and D-loop) sequences. Numbers above branches indicate posterior probability and bootstrap values, respectively. Phylogenetic relationship of Sidi Rachid population was different depending on Bayesian (A) of ML (B) topology.

Overall percentages of uncorrected-p distances ranged from 0.0 to 1.2% for *MT-CYB* and from 0.0 to 1.8% for the D-loop region (Table S2 in Supporting Information). Genetic distances between trout populations from Drâa Basin (Clade I) and the other Moroccan populations (Clade II) ranged from 0.9 to 1.2% for *MT-CYB* and from 1.1 to 1.8% for D-loop (Table S2 in Supporting Information). The genetic distances among the Moroccan populations within Clade II ranged from 0.0 to 1.0% and 0.1 to 0.8% for *MT-CYB* and D-loop, respectively.

Haplotype network analysis of *MT-CYB* supported the phylogenetic analyses (Figure 3). The analysis of all sequences obtained for *MT-CYB* showed the existence of five clearly differentiated Moroccan populations that did not share any haplotypes with the other populations: Drâa Basin, Rheraya R., Tessaout R., Sidi Rachid R. and Isli Lake. In the other Moroccan populations, most of the haplotypes were shared. Rifian populations (Farda and Kannar rivers) showed only one haplotype shared by all individuals from these basins. This haplotype, surprisingly, was also the most common one found in the Ifni and Tifnoute populations. It was also observed in the Ourika river population. On the other hand, most tributaries of the Oum er Rbia Basin were clustered in a differentiated haplogroup. Nevertheless, haplotypes of some of the tributaries of the Oum er Rbia Basin showed a closer relationship with those of other trout populations. For instance, a haplotype from Melloul were more closely related to one found in Ifni, Tifnoute and Rifian populations, or Lakhdar and Ait Nacer haplotypes were related to but not shared with those found in Sidi Hamza. The Sidi Rachid population also constituted a haplogroup related to Sidi Hamza. Tamda and Moulouya shared their two most common haplotypes. One of the museum specimens from the Beth R. (Sebou Basin) showed the same *MT-CYB* haplotype as the most frequent one found in the Sidi Hamza and Tamda populations, whereas the two museum specimens from Algeria did not shared haplotypes with any of the analyzed populations. The D-loop haplotype network analysis (Figure 4) differed slightly from the *MT-CYB* analysis for some of the relationships for the Moroccan populations excluding the Drâa Basin. The most common haplotype of Isli Lake was shared with Sidi Hamza, Lakhdar, Ait Nacer and Moulouya populations; however, the remaining haplotypes of Isli Lake were unique. In contrast with the *MT-CYB* network, Rifian populations did not share its unique haplotype with the Ifni and Tifnoute populations, and some of the Sidi Rachid haplotypes were shared with Oum er Rbia tributary populations. Ourika was closely related to some Moulouya basin individuals. Finally, Rheraya and Tessaout populations appeared as differentiated haplogroups, as was observed in the *MT-CYB* network. The relationships of the three museum specimens were the same as in the *MT-CYB* network.

**Figure 3.**
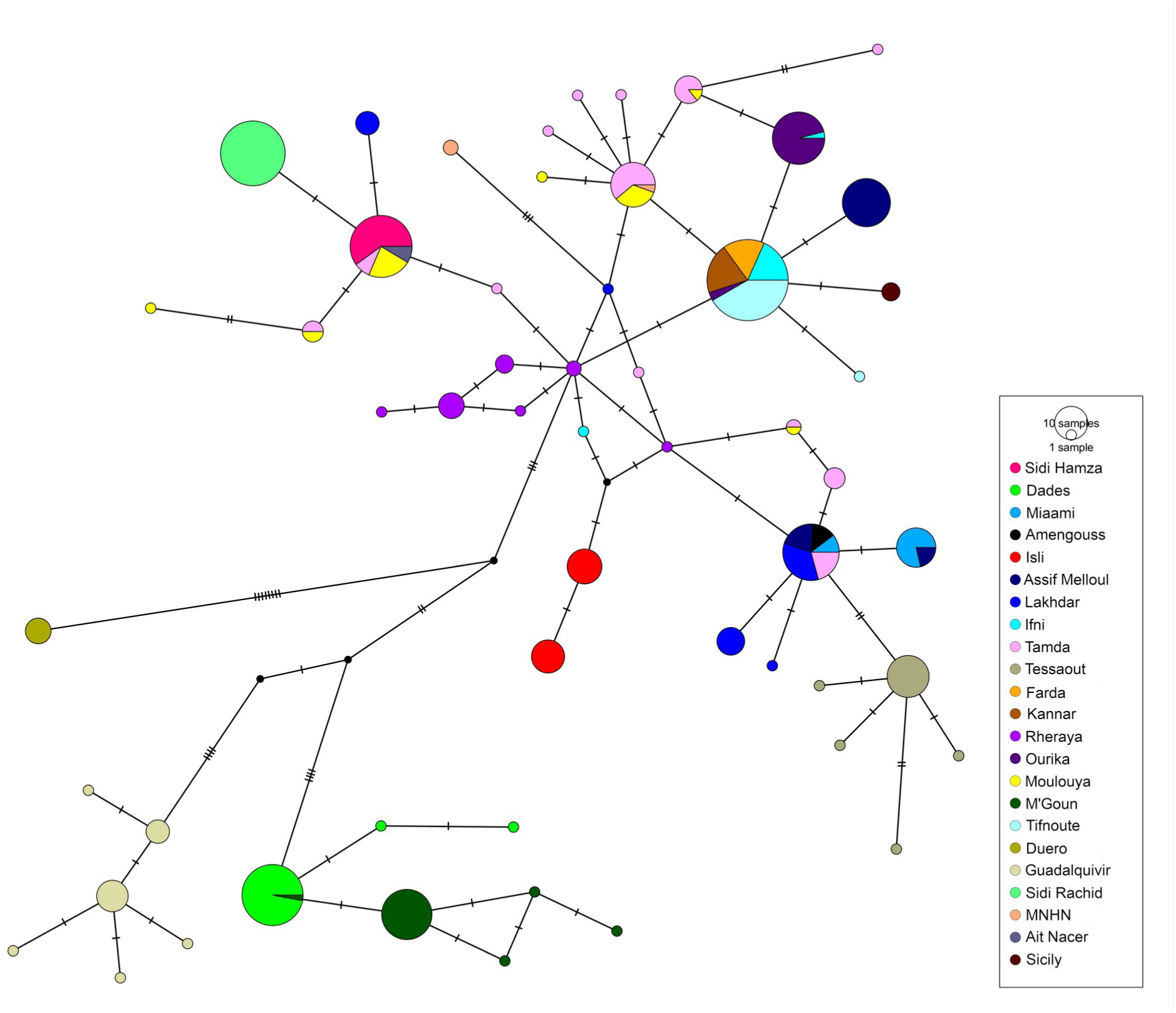
Haplotype network based on mitochondrial *MT-CYB* sequences. Dashes in branches represent the number of mutational steps.

**Figure 4.**
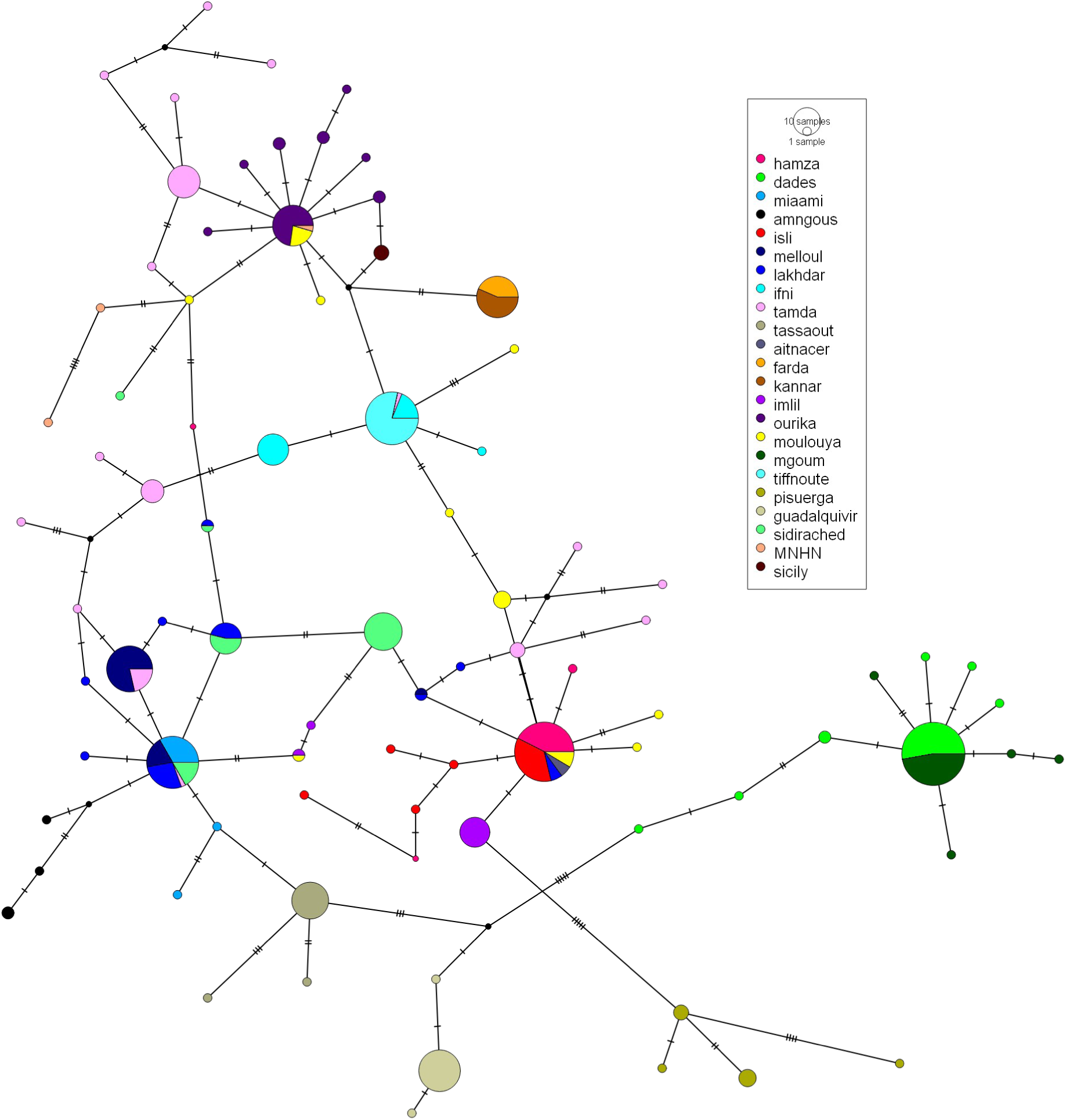
Haplotype network based on mitochondrial D-loop sequences. Dashes in branches represent the number of mutational steps.

According to divergence time estimates, Clades I, II and III diverged near the boundary of the Pliocene and the Pleistocene, ca. 5.3 Mya (3.6–6.6 95% HPD) (Figure 5). Within Clade I, the Dades and M’Goun populations (Drâa Basin) diverged from their sister lineages during the Upper Pliocene ca. 4.4 Mya (2.8–5.8 95% HPD), and from each other ca. 0.7 Mya (0.1–1.4 95% HPD) during the Pleistocene. Diversification of Moroccan popuations in Clade II started in the Upper Pliocene ca. 2.9 Mya (1.6–4.3 95% HPD) (Figure 5).

**Figure 5.**
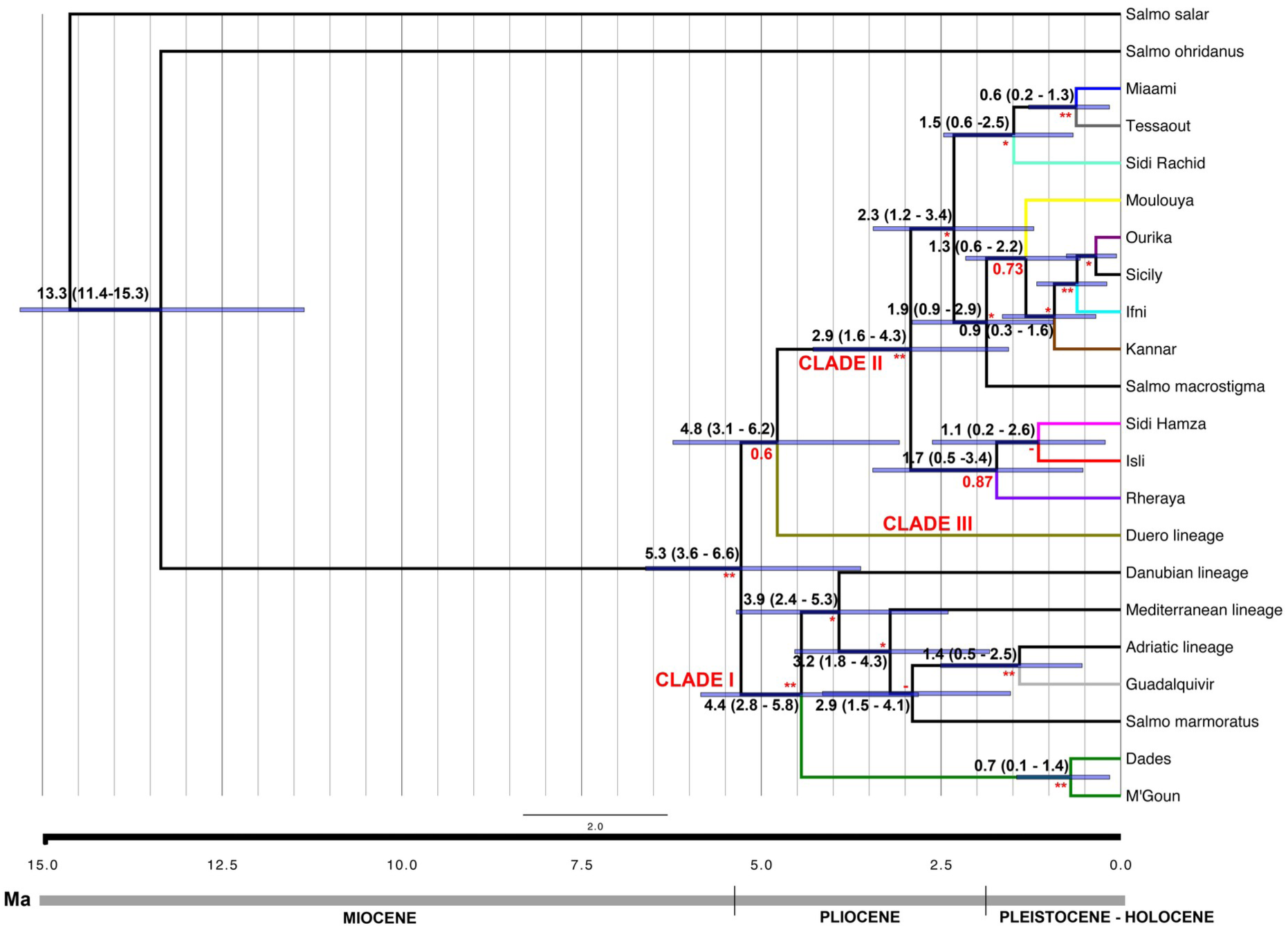
Divergence time estimations of Moroccan trout populations. Divergence time estimations and corresponding confidence intervals (HPD 95%) are indicated above branches. Posterior probability (pp) values are indicated below branches. ** pp=1; * pp>0.95.

### 3.2. Microsatellites and Mitochondrial structure analyses

The STRUCTURE analysis based on microsatellites rendered the best clustering at K=17 according to the ΔK parameter (Figure 6A). This grouping was highly congruent with the mitochondrial data, supporting Drâa Basin, Isli Lake and both tributaries of the Tensift Basin (Ourika and Rheraya) as differentiated genetic groups. The population structure within the Oum er Rbia Basin was higher than that inferred by the mitochondrial data. Tessaout constituted a well-differentiated group, while the other Oum er Rbia populations showed different alleles, although with a high level of admixture among populations (Figure 6A). This Bayesian clustering analysis also supported the mitochondrial relationship of the Rifian populations (Farda and Kannar) with those from the Ifni and Tifnoute; although Farda had low probability of assignment (Q) values due to admixture. In addition to Farda, admixture was evident in the Sidi Hamza, Moulouya, Tamda and Sidi Rachid populations and in those of the Oum er Rbia Basin. The ΔK also supported the clustering of K=7 (Figure 6A) and, hence, genetic differentiation among some of the basins. The Bayesian clustering analysis using TESS (Figure 6B) also indicated K=17 as the optimal value for both admixture models (CAR and BYM), as supported by the lowest DIC (9158.23 and 9482.29, respectively). This result was highly congruent with the STRUCTURE results (Figure 6A). TESS estimation based on the CAR admixture model recovered Ifni and Tifnoute as different populations.

**Figure 6.**
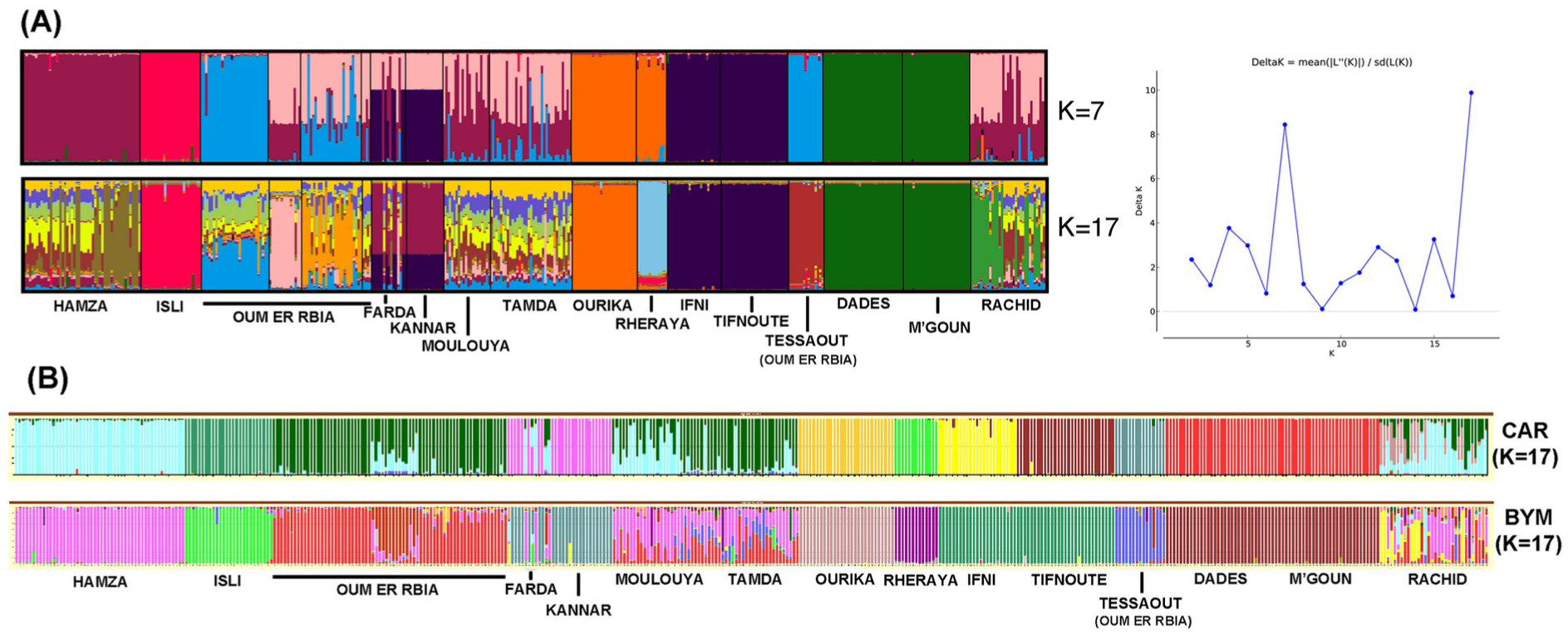
Clustering analyses of microsatellite data using STRUCTURE (A) or TESS (B). Bar plot of estimated membership of each individual in K=7 and K=17 clusters. In (A), the number of Moroccan trout populations with the highest posterior probability expressed as the ΔK is represented after the bar plot.

In the factorial correspondence analyses (FCA), the first fourth components explained 38.4% of the genetic variance. According to the FCA, four well supported genetic groups could be identified: populations from the Ifni and Tifnoute basins, the Drâa Basin (Dades and M’Goun), the Mediterranean (Farda and Kannar) and the rest of the populations (subgroup I) (Figure 7A). When only subgroup I populations were considered, populations from the Tensift Basin (Ourika and Rheraya) were differentiated from each other and from the other populations (subgroup II) (Figure 7B). Finally, when Tensift basin populations (and previous groups) were removed from the analysis, populations from Isli Lake and Tessout, a tributary of the Oum er Rbia Basin, were recovered as differentiated genetic groups, whereas Sidi Hamza, Sidi Rachid, Tamda Lake and the Mediterranean Moulouya Basin grouped with the remaining tributaries of the Oum er Rbia Basin (Figure 7C).

**Figure 7.**
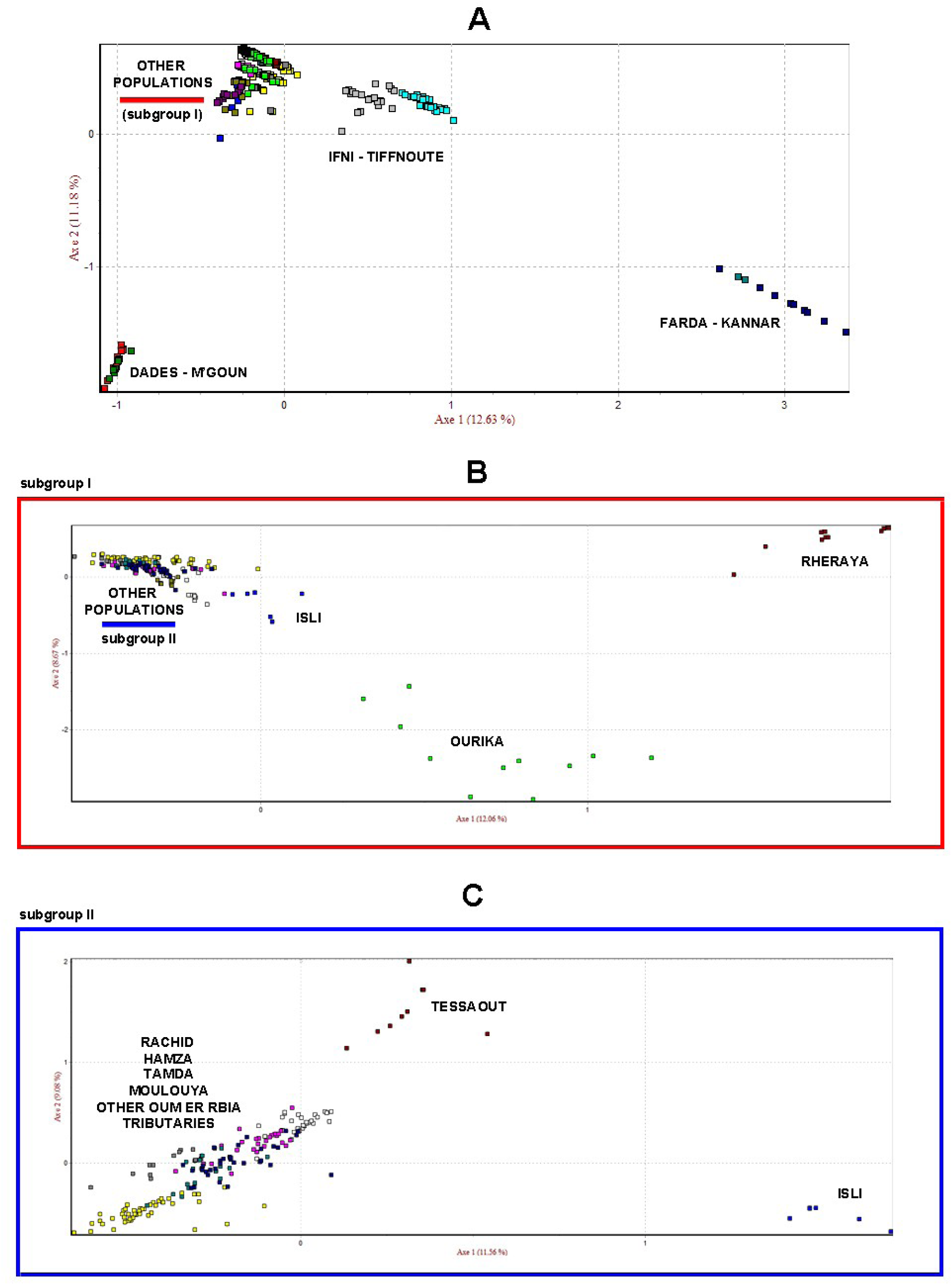
Factorial correspondence analyses (FCA) of Moroccan trout populations based of the allelic frequency of nine microsatellites. Each point represents one individual from the 16 sampling sites analyzed. The first two axes resulting from the analysis are shown. A) considering all populations. B) excluding the genetic groups of Dades-M’Goun, Farda-Kannar and Ifni-Tifnoute. C) excluding the genetic groups of Ourika and Rheraya.

All *F*_ST_ comparisons based on microsatellite data were significant (Table S3 in Supporting Information). For mitochondrial data, the Φ_ST_ pairwise comparisons among Moroccan trout populations ranged from 0.000 (between the two Rifian populations) to 1.000 (between Sidi Hamza and Farda) for *MT-CYB* and from 0.027 (Dades and M’Goun) to 1.000 (Tifnoute and Kannar) for D-loop (Table S4 in Supporting Information). The majority of comparisons were significant after Bonferroni correction. Significant levels of population differentiation were also found when the entire dataset was analyzed as one gene pool or when basins were considered as independent in the AMOVA analyses based on microsatellites and mitochondrial data (Table 2 and 3). The highest partition of genetic variance was found among basins: 58.8% for *MT-CYB* and 71.2% for D-loop. Similar to the mitochondrial analysis, the highest percentage of genetic variance (61.9%) was explained among basins when one gene pool was considered. When all basins were considered as independent, variance in all partitions was significant, with the highest percentage of genetic variance explained, again, among basins in all performed analyses.

**Table 2.**
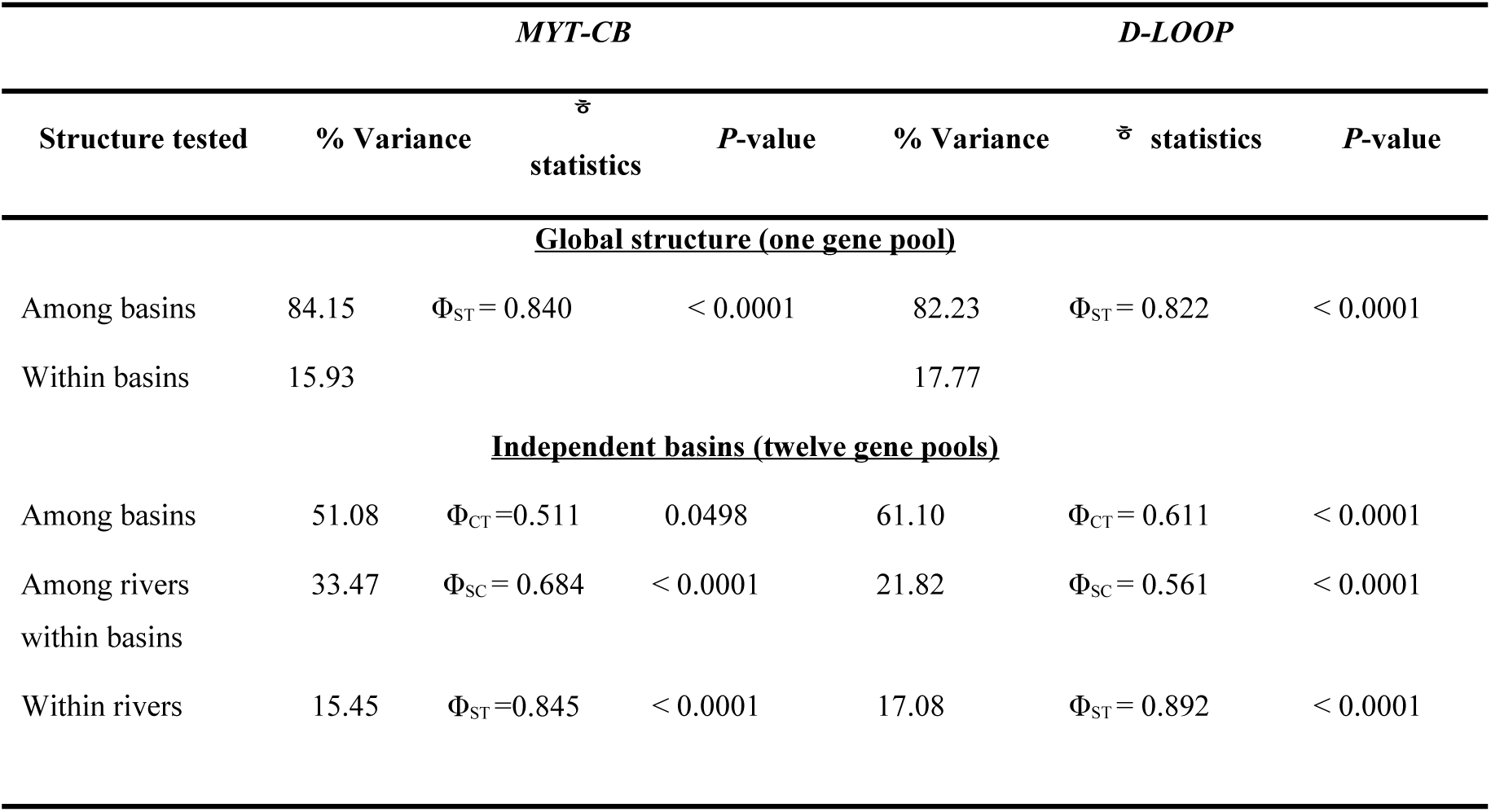
Genetic hierarchical population structure (AMOVA) for both mitochondrial markers. Analyzed basins (genetic pools): Drâa, Farda, Ifni, Isli, Kannar, Moulouya, Oum er Rbia, Sebou, Tamda, Tensift, Souss and Ziz.

**Table 3.** Genetic hierarchical population structure (AMOVA) for the nine microsatellite markers. Analyzed basins (genetic pools): Drâa, Farda, Ifni, Isli, Kannar, Moulouya, Oum er Rbia, Sebou, Tamda, Tensift, Souss and Ziz.

| Structured tests | % variance | Statistics | p-value |
| --- | --- | --- | --- |
| **<u>Global structure (one gene pool)</u>** |  |  |  |
| Among basin | 61.95 | $F_{ST} = 0.6195$ | < 0.0001 |
| Within rivers | 38.05 |  |  |
| **<u>Independent basins (twelve gene pools)</u>** |  |  |  |
| Among basins | 46.01 | $F_{CT} = 0.4601$ | < 0.0001 |
| Among rivers within<br>basins | 16.73 | $F_{SC} = 0.3098$ | < 0.0001 |
| Within rivers | 37.27 | $F_{ST} = 0.6273$ | < 0.0001 |

### 3.3. Mitochondrial and microsatellite genetic diversity analyses

Within the Moroccan populations, a total of 46 haplotypes were recognized in *MT-CYB* and 74 in D-loop. However, global haplotype diversity was very high and similar for both mitochondrial markers: 0.929 and 0.937 for *MT-CYB* and D-loop, respectively (Table S3 in Supporting Information). Global nucleotide diversity was also high (>0.5%); however, when populations were considered independently, none exceeded 0.3%, the highest value found in Tamda population (Table S5 in Supporting Information). Tamda Lake had the most haplotypes for both markers. This population also showed the highest genetic diversity values, followed by the Mediterranean Moulouya Basin. The populations from Lakhdar and Rheraya rivers also exhibited high values of mitochondrial genetic diversity. Some of the lowest genetic diversity values were found for Isli, Tifnoute, Sidi Hamza, Sidi Rachid and both Rifian populations. In fact, only one haplotype for each mitochondrial marker was found in each of the Rifian populations, as was the case for the Sidi Hamza and Sidi Rachid populations for the *MT-CYB* gene.

For microsatellites, deviation from Hardy-Weinberg Equilibrium (HWE) was not consistently observed across loci and populations except for the STR15 locus and Farda population after Bonferroni correction (adjusted α = 0.05/9 = 0.005). According to MICRO-CHECKER, null alleles were only found for STR15, accounting for its deviations from HWE. Nevertheless, the FreeNA analysis, after applying the ENA correction, indicated that potential bias on global FST calculations caused by null alleles was insignificant (Global FST = 0.627; Global FST with correction for null alleles = 0.617). Therefore, locus STR15 was included in analyses of population structure. The test for linkage disequilibrium showed a very low number of significant pairwise comparisons, indicating that all examined loci can be considered independent.

The total number of nuclear alleles detected was 91. The number of alleles per locus ranged from 3 (STR541 and STR60) to 28 (STR103), with an allelic richness of 2.85 and 8.40, respectively. Seven of the nine loci analyzed showed a total of 41 private alleles at different frequencies in the populations, from between 0.0167 and 0.0357 in Melloul, Ifni, Dades and M’Goun to 1.000 in Isli, Dades-M’Goun and Farda-Kannar. Indeed, Farda and Kannar (Rifian populations) shared private alleles at high frequencies in six of the loci. Drâa Basin tributaries also shared six private alleles in four loci. Oum er Rbia Basin tributary populations also showed private alleles, including, remarkably, four in Tessaout and three in Melloul. The Tamda lake population did not have any private alleles, and Mediterranean Moulouya Basin only had one private allele but at a low frequency (0.05). In contrast to the mtDNA data, global genetic variability was low (*H*o = 0.249) across loci and populations, with low allele numbers and heterozygosity values. Farda-Kannar, Dades-M’Goun and Tessaout populations had the lowest values of genetic diversity while the highest were found in Tamda, Moulouya and Miaami (Table S6 in Supporting Information). Allelic richness per loci ranged from 1.4 to 3.77 or from 1.19 to 2.97 when this parameter was weighted by sample size (Table S7 in Supporting Information). FIS values ranged from −0.135 in Miaami to 0.236 in M’Goun (Table S6 in Supporting Information). The significant negative value for the Miaami population indicates an excess of heterozygotes in contrast to the significant positive value, representing heterozygote deficiency, in both Farda and M’Goun.

### 3.4. Population size changes and gene flow based on mitochondrial and microsatellites markers

Based on the null hypothesis of population expansion, the neutrality tests supported deviations from the mutation-drift model in both populations of the Drâa Basin and in Tessaout (Oum er Rbia Basin) for both mitochondrial markers, and in Ourika (Tensift Basin) for D-loop (Table S7 in Supporting Information). A significant negative value of Fu’s *FS* was also estimated for the Mediterranean Moulouya Basin for D-loop. Estimates of the virtual number of migrants exchanged among populations and per generation were greater than one for some of the pairwise comparisons, particularly between the two tributaries of the Drâa Basin (*Nm* = 18) (Table S8 in Supporting Information). Positive gene flow (*Nm* > 1) was also estimated for populations from some of the tributaries of the Oum er Rbia Basin (Miaami, Melloul and Lakhdar), as well as for the Rifian populations, Ifni and Tifnoute basins. Gene flow between Tamda Lake and the other basins differed depending on the marker analyzed: a *Nm* > 1 was observed between Tamda and two tributaries of the Oum er Rbia Basin (Melloul and Lakhdar) for *MT-CYB* but not for D-loop. In contrast, for D-loop, a *Nm* >1 was found in pairwise comparisons of this lake population with those from Ifni Lake and the Rifian region. A *Nm > 1* between Tamda and Moulouya considering both markers was also estimated (Table S8 in Supporting Information).

The Wilcoxon test detected a recent bottleneck (p<0.05) in the Tifnoute population under all three mutation models analyzed for microsatellites (IAM, SSM and TPM) (Table S9 in Supporting Information). Significant values were also observed for Moulouya, Rheraya and Sidi Rachid under the IAM model. The mode-shift indicator test revealed a distortion of allele frequency distributions characteristic of a recent bottleneck for Miaami, Rheraya and Sidi Rachid. Additionally, analysis of the virtual number of migrants with the microsatellite dataset (Table S3 in Supporting Information) showed *Nm* values greater than one in pairwise comparisons of Tamda Lake with some of the Oum er Rbia tributaries (Melloul and Lakhdar), Sidi Hamza, Moulouya and Sidi Rachid. The *Nm* was also greater than one in comparisons of Sidi Rachid with Sidi Hamza, Molouya and Miaami populations, as well as between Sidi Rachid and Moulouya. The Drâa populations, Dades and M’Goun, also showed a *Nm* value greater than one.

## 4. Discussion

Understanding the evolutionary processes of species of socioeconomic interest like trout is a challenge due to the difficulty of isolating natural versus human-mediated patterns. Additionally, the Moroccan trout populations analyzed in this study are located at the southern periphery of their distribution range; thus, the evolutionary forces operating on these populations may lead to different patterns relative to those in more northern and central populations. These differences may, for instance, increase fragmentation and reduce genetic diversity, which would have important implications for the conservation of these populations and would become these peripheral populations more vulnerable to human impacts.

### 4.1. Phylogenetic and phylogeographic relationships of Moroccan trout populations

In general, five lineages have been traditionally recognized in the brown trout (Adriatic, Atlantic, Danubian, Marmoratus and Mediterranean), which, in turn, have been classified into three larger groups (Bernatchez et al., 1992; Bernatchez, 2001). Brown trout populations in Morocco, including the Mediterranean ones, have been considered as part of the large Atlantic lineage (Bernatchez, 2001; Snoj et al., 2011; Ninua et al., 2018). However, other authors have recognized a Siculo-North African lineage, as sister to the Atlantic one, and have included the Moroccan trout within it (Tougard et al., 2018). Considering only the analyzed Moroccan trout populations, two divergent evolutionary lineages have been identified in the present study, one comprising the Drâa basin populations in the southernmost river drainage within the distribution range of trout populations in Morocco. According to our results, this lineage is sister to the Danubian, Adriatic and Mediterranean lineages, *Salmo marmoratus* and the trout population from the Iberian Guadalquivir Basin. The distinctiveness of the Drâa population is in agreement with previous studies (Snoj et al., 2011; Doadrio et al., 2015; Sanz, 2017; Ninua et al., 2018; Tougard et al., 2018). The second lineage is composed of the remaining Moroccan trout populations, as well as the Algerian and Sicilian populations. These last relationships had been previously proposed in other phylogeographic studies (Berrebi et al., 2018).

### 4.2. Genetic structure of Moroccan trout has been driven by the geological and climatic history and by contemporary human-mediated processes

Analyses of concatenated mtDNA sequences and microsatellite data have revealed a genetic structure of brown trout populations in Morocco that can be mainly explained by historical factors associated with the complex palaeohydrology of North Africa. These historical factors have proven to have greatly influenced the genetic structure of other North African freshwater fishes (Doadrio, 1994, Machordom & Doadrio, 2001). However, in the case of brown trout, the degree and the temporal scale of divergence among populations are different to those of primary freshwater fishes, such as cyprinids or cobitids (Machordom & Doadrio, 2001; Casal-López & Doadrio, 2018). In addition, the patterns revealed in this study demonstrate the great impact of contemporary human-mediated processes on the genetic structure of the Moroccan populations of brown trout, and along with historical factors have been responsible for the current genetic structure found in these populations.

The Atlantic Drâa Basin is estimated to have diverged from other brown trout lineages at the Upper Pliocene, which is slightly older than previous estimations for the isolation of the Drâa lineage (Snoj et al., 2011). The High Atlas Mountains are thought to have experienced an uplifting pulse event during the Plio-Pleistocene period less than 5 Mya that has been associated with drainage reorganization (El Harfi, Guiraud & Lang, 2006; Babault et al., 2012; Boulton, Stokes & Mather, 2014). This tectonic pulse, along with the geomorphological configuration of the western Mediterranean region since the Upper Miocene–Pliocene periods (Krijgsman et al., 2018), was likely responsible for the segregation of the Drâa lineage. The complex geomorphological pattern and isolation of this region has prevented the artificial introduction of trout in the Drâa populations.

The remaining Moroccan populations indicated two different conditions. Some of the populations showed a high and recent level of geographic isolation, since the Pliocene– Pleistocene period, and no human influence due to stocking, a hypothesis supported by all of the molecular analyses. Such is the case of the Tessaout R. located on the left margin of the Oum er Rbia Basin, with a steep geomorphology, geographically distant from the remaining tributaries of the basin (Melloul, Lakhdar or Miaami), which showed a smoother landscape (Babault et al., 2012), sharing mitochondrial haplotypes and nuclear alelles among them. The two tributary populations analyzed in the Tensift Basin (Rheraya and Ourika) also demonstrated a high level of genetic differentiation and Ourika resolved as phylogenetically close to the Ifni Lake and Tifnoute R. (Souss Basin) than to Rerhaya. Ourika headwaters are very close to some tributaries of the Souss Basin, including the Tifnoute, and to the Ifni Lake. Consequently, the close phylogenetic relationship and shared haplotypes between Ourika and these populations may be explained by fluvial captures, which are known to occur all over the western High Atlas Mountains during Pleistocene (Babault et al., 2012; Boulton et al., 2014). The population from Isli Lake is genetically differentiated from the other populations. The geological origin of this lake is still under debate due to its geomorphology (rounded contour and great depth). Some authors have proposed that it originated as a consequence of a meteorite impact approximately 40,000 years ago (Ibhi, Nachit, Abia, Ait Touchnt & Vaccaro, 2013; Nachit, Ibhi & Vaccaro, 2013), while others have hypothesized a similar tectonic origin as other Moroccan lakes that is associated with karstic phenomena during the Lower–Middle Pleistocene (Chaabout, Chennaoui-Aoudjehane, Reimold, Aboulahris & Aoudjehane, 2013; Ibouh et al., 2014). The degree of genetic differentiation observed in the Isli lake population and its estimated time of divergence support its ancient origin and tectonic origin (Ibouh et al., 2014); nevertherless, hydrographical past connections (Babault et al., 2012; Ibouh et al, 2014) would explain the presence of shared haplotypes between Isli and those from some of the tributaries of the Oum er Rbia and Ziz basins.

In contrast to these highly isolated and non stocked populations, trout stocking influence is evident in other populations. Thus, the Ziz basin population (Sidi Hamza) shares haplotypes and shows some level of gene flow with populations in the Oum er Rbia, Tamda and Moulouya basins. The river source of Ziz Basin is close to those of some Oum er Rbia tributaries (Babault et al., 2012), suggesting the possibility of secondary contact between these two basins. However, given the degree of admixture human-mediated contact is a more plausible explanation for the relationship among these basins. The Ziz population has been one of the main populations used in stocking programs at the Ras el Ma hatchery over the last several decades. Therefore, the presence of its mitochondrial haplotypes in the Oum er Rbia and Mouloya basins can be attributed to an artificial process. Some Rheraya and Tamda individuals were also related to some Oum er Rbia populations, as well as the most common haplotype of Ourika R. was also found in the Moulouya basin, probably as a consequence of stocking as well (Fekhaoui, Yahyaoui, Perea & Doadrio, 2016). Another population influenced by stocking is the Sidi Rachid R. (Sebou Basin); this river flows nearest to the Ras el Ma hatchery, therefore, it has been stocked with various trout populations originating from the hatchery (El Hassen et al., 2011).

The relationship found between the Mediterranean (Farda, Kannar and Algeria) and the Ifni and Tifnoute basin populations is more challenging to explain without taking into account human-mediated processes. The presence of only one *MT-CYB* haplotype in Farda and Kannar that is shared with the distant populations of Ifni and Tifnoute calls into question the natural condition of the Rifian populations, even though several mitochondrial analyses supported the Rifian populations as distinctive groups. There is evidence that Farda R. has been stocked with trout from the Ras el Ma hatchery (Table S10 in Supporting Information; Fehkaoui et al., 2016). Hence, the possibility of Tifnoute trout serving as a source in Mediterranean Rifian populations due to stocking cannot be discarded. Nonetheless, results of the Bayesian clustering analyses based on TESS supported the Rifian populations as distinct groups. Genetic differentiation of the Algerian population further supports the native status of the three Mediterranean populations. These populations probably constitute the remnants of an ancestral trout lineage that was widely distributed in the past.

### 4.3. Genetic diversity of Moroccan trout populations is associated with historical factors and trout stocking

Overall genetic diversity values based on mitochondrial data were high, taking into account the concatenated dataset. Nevertheless, genetic diversity levels differed among individual populations and while some populations exhibit high values in others genetic diversity was extremely low. Tamda and Moulouya showed the highest levels of haplotype diversity as well as heterozygosity according to the results of the microsatellite analyses. Human-mediated origins for Tamda and Moulouya could explain the genetic sub-structure as a consequence of haplotype sharing with other basins (Sidi Hamza, Oum er Rbia and Ourika) and the high levels of diversity observed in these populations, although the Moulouya haplotypes related to those in Ourika probably constitute a natural lineage of this population. However, Moulouya was restocked two years prior to being sampled for the present study, and the high level of genetic diversity found in this population is in agreement with the use of different native trout populations to increase genetic diversity, as employed by the management stocking programs of Ras el Ma hatchery. In the case of Tamda Lake, the high genetic diversity but lack of distinctive genetic features supports the artificial origin of this population, as has been maintained by local inhabitants of the region. Hence, the high virtual number of migrants found in the Moulouya and Tamda populations may be a consequence of restocking events rather than reflecting a real gene flow process.

Lakhdar and Rheraya also displayed high levels of genetic diversity. Lakhdar showed the highest microsatellite allelic richness after weighting per sample size. Lakhdar haplotypes related to those in Sidi Hamza, a probable consequence of restocking as indicated by the clustering analyses, may explain the high mitochondrial genetic diversity uncovered in this population relative to other Oum er Rbia populations. Population expansion was not inferred for Lakhdar (non-significant neutrality tests and significant raggedness index), and stable population sizes may favor the maintenance of high genetic diversity in this population. Rheraya also does not appear to have undergone population expansion. In fact, a significant recent bottleneck was estimated to have occurred in this population on the basis of molecular analyses. Stochastic and unpredictable events associated with drastic climatic episodes during the Quaternary (Benito et al., 2015) may have influenced the genetic and historical demography of Rheraya. Given that recurrent bottlenecks can lead to the loss of genetic diversity in freshwater organisms (Allendorf, 2017; Carim, Eby, Barfoot & Boyer, 2017), the high diversity values obtained for Rheraya from the analysis of *MT-CYB* is difficult to explain without considering it a possible effect of stocking.

In contrast, highly geographically isolated and not stocked populations including Drâa Basin tributaries, Tessaout, and Isli Lake showed low levels of genetic diversity and low allelic richness. Neutrality tests supported deviations from the mutation-drift model in both populations of the Drâa Basin and in Tessaout for both mitochondrial markers. These deviations could be associated with recent population expansion, a hypothesis supported by the star-shape of the Dades-M’Goun and Tessaout haplogroups in both mitochondrial haplotype networks in which several low frequency haplotypes are connected to the most frequent one. In addition, no evidence of bottlenecks was recorded for these populations. However, the mismatch distribution (Figure S1 in Supporting Information) did not fit a population expansion model (multimodal shape), except for M’Goun (unimodal shape and significant raggedness index). Genetic drift is an evolutionary force driving loss of genetic diversity due to the random fixation along time of specific alleles, and its effect is related to effective population sizes (Kliman, Sheehy & Schulz, 2008; Frankham, Bradshaw & Brook, 2014). Given this context, genetic drift may explain the low genetic diversity found in the Drâa Basin and Tessaout populations. The geographic isolation of these three populations in rivers flowing through deep canyons in the High Atlas Mountains, along with severe climatic conditions during the Pleistocene (Hughes et al., 2011; Babault et al., 2012), could have favored small but constant effective population sizes, thereby increasing the effect of genetic drift, which would lead to decreased genetic diversity (e.g. Hare et al., 2011). Indeed, species occupying narrow altitudinal ranges, such as the trout in the Drâa Basin and Tessaout R., are particularly vulnerable to extreme climatic events (La Sorte & Jetz, 2010; Clavero et al., 2017).

Isli Lake constitutes a population that has been isolated since the Lower–Middle Pleistocene (Ibouh et al., 2014; this study). Genetic drift may have also led to the low genetic diversity observed in this population. Although Isli Lake does not appear to have experienced any bottlenecks, five of the nine microsatellites analyzed were monomorphic for this population, probably as a direct effect of inbreeding. Therefore, the bottleneck analyses should be considered with caution. In contrast, Ifni Lake showed moderately higher levels of genetic diversity relative to Isli Lake, probably due to an inflow of genotypes from ancient connections with geographically close rivers/basins such as the Tifnoute R. in the Souss Basin (Babault et al., 2012). Genetic diversity values were extremely low in Sidi Hamza (mitochondrial), Tifnoute (mitochondrial and nuclear) and both Rifian (mitochondrial and nuclear) populations. A significant bottleneck was estimated for Tifnoute based on the three microsatellite mutation models tested; however, as in Isli Lake, bottleneck events could not be inferred for Sidi Hamza and Rifian populations as most of the loci were monomorphic for these populations. However, due to the narrow geographic range of these populations, small effective population sizes would be expected.

### 4.4. Implications for conservation of Moroccan trout populations

The conservation status of Moroccan trout is, in general, poor, and some populations have been proposed to be Endangered or Critically Endangered, according to the IUCN categorization (e.g. Doadrio et al., 2015; Clavero et al., 2017). Management schemes for conservation often require an understanding of population dynamics in order to achieve effective long-term results. A central concept in biodiversity conservation is that genetic diversity is crucial to ensure the survival of species (Frankham et al. 2014). Therefore, its conservation has become an explicit goal of strategic plans such as the one implemented at the Convention on Biological Diversity (http://www.cbd.int). In fact, in conservation biology, genetic structure and genetic diversity are recognized as important criteria to consider when prioritizing populations for protection. They are also biodiversity components to be preserved according to the IUCN (Frankham et al., 2014; McGowan, Traylor-Holzer & Leus, 2017). Genetic diversity can determine factors such as species viability, resilience to environmental stressors and adaptation to changing environmental factors (Frankham et al., 2014; Rominguer et al., 2014). The preservation of genetic diversity is based on the relationship of these parameters with an organism’s potential to evolve, thus generating a background from which new variants can arise. These new variants can then potentially colonize new environments and respond adaptively to environmental changes (Duglosh, Anderson, Braasch, Cang & Gillette, 2015; Szucs, Melbourne, Tuff & Hufbauer, 2017).

The low level of genetic diversity shown here, particularly for some of the native and highly isolated populations, such as those in the Drâa Basin, Tessaout R. and Isli or Ifni lakes, is one of the main threats to Moroccan trout populations. The geographical isolation of these Moroccan trout populations, which are located in the southern periphery of the distribution range of the brown trout, also contributes to reduce genetic diversity. The isolated trout populations of the High Atlas Mountains inhabit fragmented and reduced and unstable habitats (Doadrio et al., 2015; Clavero et al., 2017). For this reason, they are more vulnerable to the effects of catastrophic events, such as flooding following an unpredictable torrential rainfall (Zkhiri, Tramblay, Hanich & Berjamy, 2017), which, in turn, can lead to changes in demographic and genetic patterns. Such changes likely have caused the low level of genetic diversity found in these isolated Moroccan trout populations along the evolutionary time. Low levels of genetic diversity increase the vulnerability and the potential extinction risk of freshwater fish populations (Faulks, Kerezsy, Unmack, Johnson & Hughes, 2017; Pavlova et al., 2017), particularly as they relate to inbreeding and reduced reproductive fitness, and may ultimately lead to a loss of evolutionary potential (Allendorf, England, Luikart, Rithcie, & Ryman, 2008). Although F_IS_ values were not significant for homozygote excess in the majority of the studied Moroccan trout populations, some were monomorphic for some of the microsatellite loci analyzed, including Isli, Kannar, Ourika, Tessaout, Dades and M’Goun. Inbreeding may be especially high in Isli Lake, where fin and snout malformations in adults are common, and the presence of juveniles is scarce (authors’ personal observation).

Due to their isolated and southern peripherial nature, the instability of their habitats and the low genetic diversity of Moroccan trout populations, the management policy of the Ras el Ma hatchery was to maintain trout stocks mixing different native populations with the aim of increasing the genetic variability of restocked populations. The main source populations kept at the hatchery are from Sidi Hamza, Sidi Rachid and some tributaries from the Oum er Rbia Basin. From this mixed pool, Moroccan native populations have been frequently reinforced in several basins (Table S10 in Supporting Information). The conservation policy executed by Moroccan environmental authorities in relation to native trout populations involves habitat improvement, introduction of new populations in regions with suitable habitats and population reinforcement (Fehkaoui et al., 2016). Overexploitation and catastrophic events that have occurred in some Moroccan regions inhabited by trout over the last 20 years have justified their reinforcement (Roman & Ait Hssane, 2014; Gaume et al., 2016; El Fels et al., 2018). These events are thought to have also reduced population sizes in trout populations in other regions of the world (George, Baldigo, Smith & Robinson, 2015; Pujolar, Vicenzi, Zane & Crivelli, 2016). Nevertheless, the current management policy of stocking trout using native Moroccan populations (i.e. translocations) to increase population size and genetic diversity in the stocked populations, which has been also implemented in other regions for the brown trout (e.g. Prodöhl, et al., 2019), may be misguided mainly as a consequence of two main reasons: the negative impact of fish stocking and the loss of local adaptations.

Trout stocking for fishing is a common practice worldwide (Petereit et al., 2018; Vera et al., 2018). In Mediterranean countries, the impact of releasing trout from hatcheries as a means to reinforce natural populations is one of the main threats for the conservation of these populations, especially in rivers in which demand for sport fishing outweighs productivity (Škraba et al., 2017; Saint-Pé et al., 2018). In Morocco, stocking of High Atlas trout populations is infrequent, supported also by the performed molecular analyses, as most populations are located in difficult to access areas, such as the tributaries of the Drâa basin or Tessaout R. Nevertheless, stocking is relatively frequent in more accessible rivers, such as some tributaries of the Oum er Rbia basin, Moulouya basin or Rheraya R. in the Tensift basin and the Tamda Lake (Fekhaoui et al., 2016). The stocking activities in these basins are responsible for the high levels of genetic diversity observed in these populations, which cannot be associated with natural evolutionary processes according to our analyses. Within the Rifian populations, a signal of introgression was found in Farda but not in Kannar, supported also by literature (Fekhaoui et al., 2016). The high proportion of shared mitochondrial haplotypes and nuclear microsatellite alleles between Sidi Hamza and Sidi Rachid with other trout populations is also difficult to explain unless one considers these two populations as the main sources of a trout stock that is repopulating the other introgressed populations. Research efforts are also hampered by stocking practices as they make it even more difficult to accurately infer the evolutionary history of salmonids (Valiquette, Perrier, Thibault & Bernatchez, 2014; but see White, Miller, Dowell, Bartron & Wagner, 2018). Morocco, a country important for trout diversity (Delling & Doadrio, 2005; Doadrio et al., 2015; Tougard et al., 2018), is not an exception, and the evolutionary history of Moroccan brown trout could be misinterpreted due to the difficulty of discerning the impacts of continuous stocking since 1957, mainly from the Ras el Ma hatchery.

The negative impacts of trout stocking may be summarized in aspects such as genetic erosion, introduction of pathogens and parasites, predation, ecological competition or even alteration of stream ecosystems (Alexides, Flecker & Kraft, 2017). Some management policies focused on stocking in alpine lakes have revealed limited genetic impact on the wild stock, even after several years of stocking (Heggenes, Roed, Hoyheim & Rosef, 2002). However, this is not the case of Moroccan trout, as admixture level in some stocked populations is high, as is suggested in this study. Moreover, several studies have demonstrated that interbreeding between farmed and wild populations may lead to loss of local adaptation to specific environmental conditions due to the introduction of “maladaptive” genotypes (Skaala et al., 2006; Bourret, O’Reilly, Carr, Berg & Bernatchez, 2011). The highly isolated Moroccan trout populations have been probably subject to local selective pressures that have led to local adaptation processes, as has been suggested to occur quite frequently in other salmonids (García de Leaniz et al., 2007; Fraser, Weir, Bernatchez, Hansen & Taylor, 2011). Differentiation of the brown trout populations in Morocco dates from the Upper Pliocene (Drâa lineage) or Pleistocene (remaining populations). In the time since their divergence, evolutionary adaptation to habitats found along the distribution range could have given rise to evolutionary trajectories linked to different environments. Therefore, artificial introductions, even if could mean an increase of genetic diversity, may negatively impact the evolutionary potential of populations and their genetic integrity due to introgressive hybridization, as has been frequently described in salmonids (Bourret et al., 2011; Sušnik, Pustovrh, Jesenšen & Snoj, 2015; Muhlfed et al., 2017). For this reason, maintaining the genetic integrity of populations such as Isli, Ifni, Drâa or Tessaout, despite their low genetic diversity, is essential. Thus, conservation measures different than those currently in effect in Morocco, and aimed at keeping local adapted populations or at creating genetic refuges, as has been proposed for the Iberian Peninsula (Araguas et al., 2017; Vera et al., 2018), should be planned, especially for those geographically isolated populations from High Atlas Mountains.

## 5. Conclusion

The native populations of brown trout in Morocco constitute a very singular entity due to their intrinsic characteristics and restricted ecological requirements of cold and well-oxygenated waters, only found in some freshwater systems of the High Atlas and Rifian mountains. These restrictive conditions limit the effective population size of some populations, as these habitats are not abundant in Moroccan mountain systems. Together with the jagged orography and geomorphology of the majority of rivers in the High Atlas, these conditions favor the geographical isolation of Moroccan trout populations. Moreover, the ever-increasing exploitation of water resources by people living in High Atlas settlements as well as the different trout stocking programs that do not take into account the genetic origin of populations have led to the poor conservation status of Moroccan trout populations.

The findings presented here also highlight the need to consider the influence of contemporary processes related to anthropogenic activity on overlapping historical evolutionary processes. The genetic analyses of trout populations from Morocco using mitochondrial and microsatellite markers have distinguished between introgressed (Lakhdar, Moulouya, Tamda, some tributaries of the Oum er Rbia basin, Sidi Rachid and Farda) and non-introgressed (Isli, Ifni, Tifnoute, Kannar, Tessaout, Ourika and Drâa Basin) populations. These analyses have revealed the low genetic diversity of the majority of native populations and a high level of inbreeding of some populations such as in Isli Lake. These analyses have also allowed evaluating the impact of the trout stocking program implemented in Morocco over the last several decades that have not stringently considered the genetic origins of populations. The high genetic diversity values of, for instance, Moulouya and Tamda populations is likely an artifact, a consequence of being a mixed population of different origins as a result of human-mediated processes. Other native populations reinforced with trout individuals from the Oum er Rbia Basin also showed high levels of genetic diversity. However, the conservation status of these highly diverse populations cannot be considered satisfactory as other stocking-related problems, such as loss of local adapted genotypes, may be affecting these populations. Hence, management programs intended to reinforce Moroccan trout populations should be revised.

## Acknowledgments

We thank P. Garzón, A. Doadrio and I. Doadrio Jr. for their help in field sampling. A. Machordom gave us helpful suggestions during microsatellite performance and I. Hortelano provide some help in laboratory procedure. Melinda Modrell helped with the English editing of this manuscript. This study was supported by the MENFPESRS and CNRST from Morocco under grant N° PPR1/2015/2 for the project “Impact des changements climatiques sur la diversité génétique des poissons des eaux douces du Maroc”. The High Commissioner for Water, Forests and Fight Against Dessertification of Morocco provided permission for fish collection.

