## Supplementary tables and figures for "Influence of historical and human factors on genetic structure and diversity patterns in peripheral populations: implications for the conservation of Moroccan trout"

**Table S1**. Best scheme partition for phylogenetic performance.

| GENE PARTITION | BASE PAIRS | EVOLUTIONARY MODEL |
| --- | --- | --- |
| D-LOOP | 1000 | TVM + I +G |
| MT-CYB 1^ST^ POSITION | 1140/3 = 380 | K81 + 1 |
| MT-CYB 2^ND^ POSITION | 1140/3 = 380 | TVM + I + G |
| MT-CYB 3^RD^ POSITION | 1140/3 = 380 | TrN + G |

**Table S2**. Uncorrected-*p* genetic distances in percentage among Moroccan trout populations based on *MT-CYB* (below diagonal) and D-loop (above diagonal). Uncorrected-*p* genetic distances within Moroccan trout populations are indicated in bold in the diagonal line for *MT-CYB* and D-loop, respectively.

|  | Hamza | Dades | Miaami | Isli | Melloul | Lakhdar | Ifni | Tamda | Farda | Kannar | Rheraya | Ourika | Moulouya | M’Goun | Tifnoute | Tessaout | Rachid | Amengouss | Ait Nacer |
| --- | --- | --- | --- | --- | --- | --- | --- | --- | --- | --- | --- | --- | --- | --- | --- | --- | --- | --- | --- |
| Hamza | **0.0/0.0** | 1.1 | 0.5 | 0.1 | 0.6 | 0.4 | 0.5 | 0.6 | 0.5 | 0.5 | 0.2 | 0.7 | 0.3 | 1.1 | 0.4 | 0.5 | 0.3 | 0.8 | 0.0 |
| Dades | 1.0 | **0.0/0.1** | 1.2 | 1.1 | 1.3 | 1.2 | 1.6 | 1.6 | 1.6 | 1.6 | 1.2 | 1.7 | 1.4 | 0.1 | 1.5 | 1.1 | 1.3 | 1.4 | 1.1 |
| Miaami | 0.4 | 1.0 | **0.0/0.1** | 0.5 | 0.1 | 0.2 | 0.6 | 0.6 | 0.6 | 0.6 | 0.5 | 0.8 | 0.6 | 1.2 | 0.5 | 0.2 | 0.2 | 0.3 | 0.5 |
| Isli | 0.5 | 1.1 | 0.4 | **0.0/0.1** | 06 | 0.4 | 0.5 | 0.7 | 0.6 | 0.6 | 0.2 | 0.7 | 0.4 | 1.2 | 0.5 | 0.6 | 0.5 | 0.9 | 0.1 |
| Melloul | 0.4 | 1.0 | 0.3 | 0.4 | **0.2/0.1** | 0.2 | 0.6 | 0.5 | 0.7 | 0.7 | 0.6 | 0.8 | 0.7 | 1.3 | 0.6 | 0.3 | 0.3 | 0.4 | 0.6 |
| Lakhdar | 0.3 | 1.0 | 0.2 | 0.4 | 0.3 | **0.2/** | 0.5 | 0.6 | 0.6 | 0.6 | 0.5 | 0.7 | 0.5 | 1.3 | 0.5 | 0.3 | 0.2 | 0.5 | 0.4 |
| Ifni | 0.2 | 0.9 | 0.4 | 0.4 | 0.2 | 0.3 | **0.2/0.1** | 0.4 | 0.4 | 0.4 | 0.6 | 0.3 | 0.4 | 1.6 | 0.1 | 0.8 | 0.5 | 0.9 | 0.5 |
| Tamda | 0.3 | 1.0 | 0.4 | 0.4 | 0.3 | 0.3 | 0.3 | **0.3/0.5** | 0.6 | 0.6 | 0.6 | 0.5 | 0.6 | 1.6 | 0.4 | 0.8 | 0.6 | 0.9 | 0.6 |
| Farda | 0.3 | 0.9 | 0.3 | 0.4 | 0.1 | 0.3 | 0.1 | 0.2 | **0.0/0.0** | 0.0 | 0.4 | 0.3 | 0.4 | 1.6 | 0.3 | 0.8 | 0.5 | 0.9 | 0.5 |
| Kannar | 0.3 | 0.9 | 0.3 | 0.4 | 0.1 | 0.3 | 0.1 | 0.2 | 0.0 | **0.0/0.0** | 0.4 | 0.3 | 0.4 | 1.6 | 0.3 | 0.8 | 0.5 | 0.9 | 0.5 |
| Rheraya | 0.3 | 0.9 | 0.4 | 0.4 | 0.3 | 0.3 | 0.3 | 0.3 | 0.2 | 0.2 | **0.1/0.1** | 0.6 | 0.4 | 1.3 | 0.5 | 0.6 | 0.4 | 0.9 | 0.2 |
| Ourika | 0.3 | 1.0 | 0.4 | 0.4 | 0.2 | 0.4 | 0.2 | 0.2 | 0.1 | 0.1 | 0.3 | **0.0/0.1** | 0.4 | 1.8 | 0.2 | 1.0 | 0.7 | 1.1 | 0.6 |
| Moulouya | 0.2 | 1.0 | 0.4 | 0.5 | 0.3 | 0.4 | 0.2 | 0.3 | 0.2 | 0.2 | 0.3 | 0.3 | **0.2/0.4** | 1.4 | 0.3 | 0.7 | 0.5 | 0.9 | 0.3 |
| M’Goun | 1.1 | 0.1 | 1.1 | 1.2 | 1.1 | 1.1 | 1.0 | 1.1 | 1.0 | 1.0 | 1.0 | 1.1 | 1.1 | **0.0/0.0** | 1.5 | 1.2 | 1.3 | 1.5 | 1.1 |
| Tifnoute | 0.3 | 0.9 | 0.3 | 0.4 | 0.2 | 0.3 | 0.1 | 0.2 | 0.0 | 0.0 | 0.2 | 0.1 | 0.2 | 1.0 | **0.0/0.0** | 0.7 | 0.4 | 0.8 | 0.4 |
| Tessaout | 0.5 | 1.2 | 0.3 | 0.5 | 0.5 | 0.3 | 0.5 | 0.5 | 0.5 | 0.5 | 0.5 | 0.5 | 0.6 | 1.3 | 0.5 | **0.0/0.1** | **0.4** | **0.6** | **0.5** |
| Rachid | 0.1 | 1.1 | 0.5 | 0.6 | 0.3 | 0.4 | 0.3 | 0.4 | 0.4 | 0.4 | 0.4 | 0.4 | 0.3 | 1.1 | 0.4 | 0.6 | **0.2/0.0** | **0.5** | 0.1 |
| Amengouss | 0.4 | 1.0 | 0.1 | 0.3 | 0.3 | 0.1 | 0.3 | 0.3 | 0.3 | 0.3 | 0.3 | 0.3 | 0.4 | 1.1 | 0.3 | 0.2 | 0.4 | **0.2/0.0** | **0.8** |
| Ait Nacer | 0.0 | 1.0 | 0.4 | 0.5 | 0.4 | 0.3 | 0.2 | 0.3 | 0.3 | 0.3 | 0.3 | 0.3 | 0.2 | 1.1 | 0.3 | 0.5 | **0.1** | 0.4 | **0.0/0.0** |

**Table S3**: Pairwise estimates of *F_ST_* for the microsatellite dataset (below diagonal) and virtual number of migrants (*Nm*) exchanged between populations (above diagonal), considering M=2N (diploid markers). All *F_ST_* comparisons between sample pairs were significant after Bonferroni correction. *Nm* > 1 in bold.

|  | Hamza | Isli | Melloul | Miaami | Lakhdar | Farda | Kannar | Mouloya | Tamda | Ourika | Rheraya | Ifni | Tifnoute | Tessaout | Dades | M’Goun | Rachid |
| --- | --- | --- | --- | --- | --- | --- | --- | --- | --- | --- | --- | --- | --- | --- | --- | --- | --- |
| Hamza | - | 0.21 | 0.42 | **1.19** | 0.68 | 0.27 | 0.10 | **2.63** | **1.13** | 0.17 | 0.35 | 0.22 | 0.36 | 0.31 | 0.10 | 0.10 | **1.10** |
| Isli | 0.54 | - | 0.18 | 0.12 | 0.28 | 0.07 | 0.01 | 0.22 | 0.32 | 0.03 | 0.09 | 0.06 | 0.07 | 0.07 | 0.02 | 0.02 | 0.23 |
| Melloul | 0.37 | 0.57 | - | 0.36 | **2.71** | 0.15 | 0.05 | **1.05** | **1.4** | 0.10 | 0.26 | 0.11 | 0.10 | 0.36 | 0.05 | 0.05 | 0.66 |
| Miaami | 0.17 | 0.67 | 0.40 | - | 0.70 | 0.30 | 0.08 | **1.33** | 0.83 | 0.12 | 0.31 | 0.20 | 0.15 | 0.23 | 0.08 | 0.07 | 0.89 |
| Lakhdar | 0.26 | 0.47 | 0.08 | 0.26 | - | 0.24 | 0.08 | **2.00** | **2.39** | 0.16 | 0.39 | 0.17 | 0.15 | 0.53 | 0.09 | 0.09 | **1.13** |
| Farda | 0.48 | 0.77 | 0.62 | 0.45 | 0.51 | - | 0.82 | 0.29 | 0.25 | 0.08 | 0.18 | 0.25 | 0.25 | 0.14 | 0.07 | 0.07 | 0.34 |
| Kannar | 0.71 | 0.95 | 0.83 | 0.75 | 0.74 | 0.23 | - | 0.09 | 0.09 | 0.02 | 0.05 | 0.10 | 0.10 | 0.03 | 0.02 | 0.02 | 0.12 |
| Moulouya | 0.08 | 0.53 | 0.19 | 0.16 | 0.11 | 0.46 | 0.74 | - | **3.44** | 0.15 | 0.38 | 0.20 | 0.17 | 0.42 | 0.08 | 0.08 | **1.54** |
| Tamda | 0.18 | 0.44 | 0.15 | 0.23 | 0.09 | 0.51 | 0.73 | 0.07 | - | 0.17 | 0.43 | 0.19 | 0.17 | 0.66 | 0.08 | 0.08 | **1.08** |
| Ourika | 0.59 | 0.86 | 0.70 | 0.68 | 0.60 | 0.74 | 0.90 | 0.63 | 0.59 | - | 0.14 | 0.09 | 0.07 | 0.06 | 0.03 | 0.03 | 0.02 |
| Rheraya | 0.41 | 0.73 | 0.48 | 0.44 | 0.39 | 0.57 | 0.82 | 0.39 | 0.37 | 0.64 | - | 0.15 | 0.12 | 0.20 | 0.06 | 0.05 | 0.54 |
| Ifni | 0.53 | 0.79 | 0.69 | 0.55 | 0.60 | 0.50 | 0.72 | 0.55 | 0.57 | 0.73 | 0.63 | - | 0.67 | 0.10 | 0.07 | 0.06 | 0.25 |
| Tifnoute | 0.58 | 0.78 | 0.70 | 0.62 | 0.62 | 0.50 | 0.70 | 0.59 | 0.59 | 0.78 | 0.67 | 0.27 | - | 0.10 | 0.06 | 0.06 | 0.22 |
| Tessaout | 0.44 | 0.78 | 0.41 | 0.51 | 0.32 | 0.64 | 0.90 | 0.37 | 0.27 | 0.80 | 0.55 | 0.70 | 0.71 | - | 0.03 | 0.03 | 0.37 |
| Dades | 0.71 | 0.92 | 0.82 | 0.75 | 0.73 | 0.77 | 0.91 | 0.75 | 0.72 | 0.89 | 0.80 | 0.78 | 0.81 | 0.88 | - | **1.71** | 0.12 |
| M’Goun | 0.71 | 0.92 | 0.82 | 0.77 | 0.74 | 0.78 | 0.91 | 0.76 | 0.72 | 0.89 | 0.81 | 0.79 | 0.81 | 0.88 | 0.13 | - | 0.12 |
| Rachid | 0.18 | 0.52 | 0.27 | 0.22 | 0.18 | 0.42 | 0.67 | 0.14 | 0.18 | 0.50 | 0.31 | 0.50 | 0.53 | 0.40 | 0.67 | 0.67 | - |

**Table S4.** Φ_ST_ pairwise comparisons among Moroccan trout populations: *MT-CYB* (under diagonal) and D-loop (above diagonal). In bold are represented the only non- significant pairwise comparisons after Bonferroni correction (p<0.0001).

|  | Hamza | Dades | M’Goun | Miaami | Melloul | Lakhdar | Tessaout | Isli | Ifni | Tifnoute | Tamda | Farda | Kannar | Rheraya | Ourika | Moulouya | Rachid |
| --- | --- | --- | --- | --- | --- | --- | --- | --- | --- | --- | --- | --- | --- | --- | --- | --- | --- |
| Hamza | - | 0.96 | 0.98 | 0.96 | 0.93 | 0.73 | 0.96 | **0.12** | 0.94 | 0.99 | 0.59 | 0.99 | 0.99 | 0.84 | 0.94 | 0.39 | 0.72 |
| Dades | 0.99 | - | 0.03 | 0.95 | 0.95 | 0.89 | 0.95 | 0.93 | 0.96 | 0.98 | 0.82 | 0.97 | 0.97 | 0.94 | 0.96 | 0.87 | 0.91 |
| M’Goun | 0.99 | 0.79 | - | 0.97 | 0.96 | 0.90 | 0.96 | 0.94 | 0.98 | 0.99 | 0.82 | 0.98 | 0.98 | 0.96 | 0.97 | 0.87 | 0.92 |
| Miaami | 0.98 | 0.98 | 0.97 | - | 0.42 | 0.16 | 0.77 | 0.84 | 0.91 | 0.97 | 0.48 | 0.95 | 0.96 | 0.87 | 0.91 | 0.60 | 0.50 |
| Melloul | 0.79 | 0.91 | 0.90 | 0.63 | - | 0.33 | 0.80 | 0.86 | 0.89 | 0.94 | 0.46 | 0.92 | 0.92 | 0.87 | 0.91 | 0.70 | 0.59 |
| Lakhdar | 0.73 | 0.91 | 0.89 | 0.37 | 0.42 | - | 0.56 | 0.63 | 0.71 | 0.76 | 0.39 | 0.72 | 0.73 | 0.62 | 0.77 | 0.43 | 0.18 |
| Tessaout | 0.97 | 0.98 | 0.97 | 0.86 | 0.74 | 0.60 | - | 0.86 | 0.93 | 0.97 | 0.60 | 0.96 | 0.96 | 0.89 | 0.93 | 0.69 | 0.74 |
| Isli | 0.96 | 0.97 | 0.97 | 0.90 | 0.74 | 0.68 | 0.91 | - | 0.84 | 0.89 | 0.55 | 0.85 | 0.87 | 0.59 | 0.86 | 0.30 | 0.65 |
| Ifni | 0.76 | 0.93 | 0.92 | 0.75 | 0.19 | 0.44 | 0.81 | 0.78 | - | 0.60 | 0.23 | 0.89 | 0.90 | 0.87 | 0.77 | 0.46 | 0.76 |
| Tifnoute | 0.99 | 0.99 | 0.98 | 0.95 | 0.41 | 0.66 | 0.95 | 0.94 | 0.23 | - | 0.39 | 1.00 | 1.00 | 0.94 | 0.84 | 0.50 | 0.81 |
| Tamda | 0.60 | 0.85 | 0.84 | 0.50 | 0.26 | 0.28 | 0.62 | 0.59 | 0.17 | 0.30 | - | 0.43 | 0.45 | 0.47 | 0.38 | 0.27 | 0.47 |
| Farda | 1.00 | 0.99 | 0.98 | 0.95 | 0.31 | 0.56 | 0.94 | 0.92 | **0.12** | **-0.04** | 0.22 | - | **0.00** | 0.87 | 0.83 | 0.46 | 0.78 |
| Kannar | 1.00 | 0.99 | 0.98 | 0.95 | 0.33 | 0.58 | 0.94 | 0.92 | **0.14** | **-0.03** | 0.24 | 0.00 | - | 0.88 | 0.84 | 0.49 | 0.79 |
| Rheraya | 0.87 | 0.94 | 0.93 | 0.78 | 0.52 | 0.49 | 0.83 | 0.79 | 0.45 | 0.75 | 0.35 | 0.65 | 0.67 | - | 0.84 | 0.32 | 0.66 |
| Ourika | 0.97 | 0.98 | 0.98 | 0.95 | 0.58 | 0.72 | 0.95 | 0.92 | 0.53 | 0.85 | 0.34 | 0.84 | 0.84 | 0.75 | - | 0.49 | 0.81 |
| Moulouya | 0.47 | 0.90 | 0.89 | 0.67 | 0.40 | 0.40 | 0.75 | 0.72 | 0.17 | 0.49 | **0.12** | 0.37 | 0.39 | 0.42 | 0.55 | - | 0.46 |
| Rachid | 0.00 | 0.99 | 0.98 | 0.95 | 0.33 | 0.58 | 0.94 | 0.92 | **0.14** | **0.00** | 0.23 | **0.00** | **0.00** | 0.72 | 0.85 | 0.39 | - |

**Table S5**. Genetic diversity statistics estimated for Moroccan trout populations based on the analysis of two mitochondrial markers. N = number of individuals; h = number of haplotypes; H_D_ = haplotype diversity; π = nucleotide diversity; S = number of polymorphic sites; k = number of pairwise differences.

| POPULATION | N | h | H_D_ (SD) | π (SD) | S | k |
| --- | --- | --- | --- | --- | --- | --- |
| *MT-CYB* | | | | | | |
| Sidi Hamza | 40 | 1 | 0.000 (0.000) | 0.000 (0.000) | 0 | 0.000 |
| Miaami | 16 | 2 | 0.325 (0.015) | 0.00028 (0.0001) | 1 | 0.325 |
| Melloul | 30 | 3 | 0.476 (0.053) | 0.0017 (0.0003) | 5 | 1.924 |
| Lakhdar | 24 | 5 | 0.725 (0.003) | 0.002 (0.0004) | 8 | 2.413 |
| Tessaout | 20 | 5 | 0.368 (0.135) | 0.0004 (0.0002) | 5 | 0.500 |
| Isli | 24 | 2 | 0.052 (0.034) | 0.0004 (0.00003) | 1 | 0.518 |
| Ifni | 17 | 4 | 0.551 (0.116) | 0.002 (0.0004) | 6 | 1.824 |
| Tifnoute | 26 | 2 | 0.077 (0.070) | 0.00007 (0.00006) | 1 | 0.077 |
| Farda | 10 | 1 | 0.000 (0.000) | 0.000 (0.000) | 0 | 0.000 |
| Kannar | 12 | 1 | 0.000 (0.000) | 0.000 (0.000) | 0 | 0.000 |
| Tamda | 40 | 13 | 0.871 (0.031) | 0.003 (0.0002) | 12 | 3.081 |
| Moulouya | 20 | 7 | 0.768 (0.069) | 0.002 (0.0002) | 8 | 2.763 |
| Ourika | 28 | 2 | 0.198 (0.092) | 0.0002 (0.00008) | 1 | 0.198 |
| Rheraya | 14 | 7 | 0.846 (0.074) | 0.001 (0.0002) | 5 | 1.747 |
| Dades | 35 | 3 | 0.113 (0.072) | 0.00015 (0.0001) | 2 | 0.168 |
| M’Goun | 27 | 5 | 0.279 (0.112) | 0.0003 (0.0001) | 3 | 0.359 |
| Sidi Rachid | 34 | 1 | 0.000 (0.000) | 0.000 (0.000) | 0 | 0.000 |
| Total | 424 | 46 | 0.929 (0.004) | 0.00486 (0.00018) | 45 | 5.551 |
| D-loop | | | | | | |
| Sidi Hamza | 23 | 2 | 0.087 (0078) | 0.00009 (0.00008) | 1 | 0.087 |
| Miaami | 15 | 3 | 0.257 (0.125) | 0.0005 (0.0003) | 3 | 0.514 |
| Melloul | 32 | 3 | 0.462 (0.074) | 0.0007 (0.0003) | 5 | 0.694 |
| Lakhdar | 26 | 10 | 0.806 (0.060) | 0.0022 (0.0004) | 8 | 2.182 |
| Tessaout | 20 | 3 | 0.195 (0.115) | 0.0005 (0.0003) | 5 | 0.500 |
| Isli | 23 | 6 | 0.395 (0.128) | 0.0009 (0.0003) | 5 | 0.941 |
| Ifni | 21 | 3 | 0.529 (0.079) | 0.0006 (0.0001) | 2 | 0.590 |
| Tifnoute | 29 | 1 | 0.000 (0.000) | 0.000 (0.000) | 0 | 0.000 |
| Farda | 10 | 1 | 0.000 (0.000) | 0.000 (0.000) | 0 | 0.000 |
| Kannar | 13 | 1 | 0.000 (0.000) | 0.000 (0.000) | 0 | 0.000 |
| Tamda | 43 | 17 | 0.856 (0.039) | 0.0048 (0.0004) | 19 | 4.731 |
| Moulouya | 20 | 10 | 0.884 (0.045) | 0.0038 (0.0004) | 13 | 3.800 |
| Ourika | 26 | 8 | 0.622 (0.107) | 0.0008 (0.0002) | 7 | 0.815 |
| Rheraya | 14 | 3 | 0.275 (0.148) | 0.00094 (0.0005) | 4 | 0.934 |
| Dades | 35 | 7 | 0.363 (0.104) | 0.0007 (0.0002) | 7 | 0.659 |
| M’Goun | 29 | 5 | 0.261 (0.106) | 0.00041 (0.00019) | 5 | 0.409 |
| Sidi Rachid | 34 | 5 | 0.631 (0.069) | 0.00155 (0.00021) | 6 | 1.538 |
| Total | 420 | 74 | 0.937 (0.004) | 0.007 (0.00022) | 77 | 6.951 |

**Table S6.** Genetic diversity statistics of the nine microsatellite loci genotyped in the analyzed *Salmo trutta* populations. N = population size; N_A_ = total number of alleles per locus and mean number of alleles per population; N_AR_ = mean allelic richness standardized to the smallest sample size using the rarefaction method of FSTAT 2.9.3 (Goudet, 2001) per locus and population; P_A_ = number of private alleles (*private alleles in the Rifian populations Farda – Kannar; **private alleles in Drâa Basin); H_O_ = observed heterozygosity; H_E_ = expected heterozygosity; F_IS_ = Wright’s statistics per locus and population. Significant values of F_IS_ are indicated in bold.

| POPULATION | N | N_A_ | N_AR_ | P_A_ | H_O_ | H_E_ | F_IS_ |
| --- | --- | --- | --- | --- | --- | --- | --- |
| Sidi Hamza | 50 | 3.000 | 2.204 | - | 0.335 | 0.322 | -0.041 |
| Isli | 26 | 1.444 | 1.191 | 2 | 0.230 | 0.037 | 0.197 |
| Melloul | 29 | 3.000 | 2.045 | 3 | 0.237 | 0.239 | 0.008 |
| Miaami | 14 | 2.667 | 2.318 | - | 0.427 | 0.378 | **-0.135** |
| Lakhdar | 26 | 3.444 | 2.969 | 2 | 0.326 | 0.355 | 0.081 |
| Farda | 14 | 3.000 | 2.523 | 6* | 0.111 | 0.433 | **0.751** |
| Kannar | 16 | 1.444 | 1.292 | 6* | 0.079 | 0.077 | -0.024 |
| Moulouya | 20 | 3.111 | 2.511 | 1 | 0.416 | 0.404 | -0.031 |
| Tamda | 34 | 3.556 | 2.631 | - | 0.392 | 0.386 | -0.018 |
| Ourika | 28 | 1.889 | 1.571 | 7 | 0.131 | 0.118 | -0.108 |
| Rheraya | 13 | 1.778 | 1.717 | 3 | 0.239 | 0.241 | 0.006 |
| Ifni | 23 | 3.000 | 2.206 | 2 | 0.335 | 0.305 | -0.102 |
| Tifnoute | 29 | 2.111 | 1.933 | 2 | 0.318 | 0.305 | -0.044 |
| Tessaout | 15 | 1.667 | 1.405 | 4 | 0.118 | 0.106 | -0.120 |
| Dades | 34 | 1.667 | 1.481 | 1 +6** | 0.140 | 0.129 | -0.089 |
| M’Goun | 28 | 1.556 | 1.337 | 6** | 0.072 | 0.093 | **0.236** |
| Sidi Rachid | 25 | 3.778 | 2.951 | 2 | 0.517 | 0.489 | **-0.038** |

**Table S7**. Demographic characteristics of each population based on the analysis of mitochondrial *MT-CYB* and D-loop markers. Fs (Fu’s Fs test); D (Tajima’s D test); R2 (Ramos-Osins and Rozas test); r (raggedness index); *τ* (statistic tau for population expansion). P-values are indicated in parentheses; significant values are indicated in bold. Null hypothesis = population growth. – indicates that no polymorphisms were found in the population.

| POPULATION | Fs (*p*-value) | D (*p*-value) | R2 (*p*-value) | *r* | *τ* | Fs (*p*-value) | D (*p*-value) | R2 (*p*-value) | *r* | *τ* |
| --- | --- | --- | --- | --- | --- | --- | --- | --- | --- | --- |
|  | ***MT-CYB*** | | | | | ***D-LOOP*** | | | | |
| Hamza | - | - | - | - | - | **-0.993 (0.005)** | -1.161 (0.144) | 0.204 (0.989) | 0.690 (1.000) | 0.087 |
| Dades | **-1.857 (0.016)** | -1.281 (0.070) | 0.088 (0.185) | 0.692 (0.790) | 0.000 | **-4.092 (0.001)** | **-1.746 (0.017)** | **0.055 (0.000)** | 0.211 (0.492) | 0.000 |
| M’Goun | **-3.599 (0.001)** | -1.307 (0.074) | 0.107 (0.224) | 0.290 (0.583) | 0.165 | **-3.122 (0.003)** | **-1.866 (0.009)** | 0.087 (0.111) | **0.390 (0.023)** | 0.010 |
| Miaami | 0.551 (0.399) | 0.156 (0.758) | 0.162 (0.337) | 0.228 (0.450) | 0.325 | -0.379 (0.248) | -1.316 (0.074) | 0.171 (0.230) | 0.425 (0.718) | 0.000 |
| Melloul | 3.772 (0.944) | 1.431 (0.932) | 0.192 (0.941) | 0.360 (0.895) | 0.287 | 0.790 (0.651) | -1.185 (0.106) | 0.146 (0.576) | **0.182 (0.026)** | 0.792 |
| Lakhdar | 1.562 (0.836) | 0.405 (0.681) | 0.144 (0.649) | 0.111 (0.591) | 0.681 | -2.785 (0.069) | 0.127 (0.608) | 0.134 (0.556) | **0.065 (0.030)** | 1.071 |
| Tessaout | **-2.991 (0.001)** | **-1.974 (0.004)** | 0.107 (0.131) | 0.174 (0.281) | 0.282 | -0.213 (0.294) | **-1.794 (0.006)** | 0.122 (0.663) | 0.038 (0.024) | 3.698 |
| Isli | 1.572 (0.745) | 1.573 (0.951) | 0.259 (0.990) | 0.270 (0.619) | 0.518 | -1.264 (0.177) | 0.896 (0.220) | 0.112 (0.217) | 0.288 (0.022) | 0.030 |
| Ifni | 1.266 (0.768) | 0.092 (0.577) | 0.142 (0.443) | 0.365 (0.876) | 0.508 | 0.142 (0.478) | 0.143 (0.666) | 0.163 (0.561) | **0.167 (0.027)** | 0.590 |
| Tifnoute | -1.093 (0.064) | -1.155 (0.144) | 0.192 (0.930) | 0.722 (0.256) | 0.077 | - | - | - | - | - |
| Farda | - | - | - | - | - | - | - | - | - | - |
| Kannar | - | - | - | - | - | - | - | - | - | - |
| Rheraya | -2.502 (0.031) | 0.385 (0.711) | 0.149 (0.317) | 0.121 (0.305) | 1.253 | 0.711 (0.639) | -0.848 (0.234) | **0.118 (0.041)** | **0.545 (0.021)** | 1.236 |
| Ourika | 0.097 (0.248) | -0.363 (0.277) | 0.099 (0.160) | 0.403 (0.591) | 0.198 | **-5.254 (0.0001)** | **-1.700 (0.025)** | **0.063 (<0.001)** | 0.111 (0.159) | 0.815 |
| Moulouya | -0.236 (0.451) | 0.759 (0.810) | 0.160 (0.748) | 0.161 (0.723) | 2.882 | **-1.366 (0.003)** | 0.143 (0.594) | 0.130 (0.519) | **0.061 (0.026)** | 3.873 |
| Tamda | 0.283 (0.682) | -2.959 (0.099) | 0.123 (0.624) | 0.099 (0.631) | 2.306 | -3.347 (0.109) | 0.251 (0.666) | 0.122 (0.688) | 0.038 (0.327) | 2.836 |
| Rachid | - | - | - | - | - | 0.692 (0.456) | 0.133 (0.10) | 0.131 (0.629) | 0.143 (0.491) | 3.708 |

**Table S8**. Gene flow estimated as the virtual number of migrants contributing alleles each generation (*Nm*) for *MT-CYB* (below diagonal) and D-loop (above diagonal). *Nm=M* considering haploid markers*.* Nm>1 are indicated in bold.

|  | Hamza | Dades | M’Goun | Miaami | Melloul | Lakhdar | Tessaout | Isli | Ifni | Tifnoute | Tamda | Farda | Kannar | Rheraya | Ourika | Moulouya | Rachid |
| --- | --- | --- | --- | --- | --- | --- | --- | --- | --- | --- | --- | --- | --- | --- | --- | --- | --- |
| Hamza | - | 0.02 | 0.01 | 0.03 | 0.04 | 0.21 | 0.03 | **5.00** | 0.04 | 0.004 | 0.40 | 0.006 | 0.006 | 0.19 | 0.04 | 0.81 | 0.20 |
| Dades | 0.003 | - | **18.0** | 0.03 | 0.04 | 0.06 | 0.03 | 0.04 | 0.02 | 0.01 | 0.11 | 0.02 | 0.01 | 0.03 | 0.02 | 0.08 | 0.05 |
| M’Goun | 0.01 | 0.13 | - | 0.02 | 0.02 | 0.05 | 0.02 | 0.03 | 0.01 | 0.01 | 0.11 | 0.01 | 0.01 | 0.02 | 0.02 | 0.07 | 0.04 |
| Miaami | 0.01 | 0.01 | 0.01 | - | 0.69 | **2.49** | 0.14 | 0.09 | 0.05 | 0.02 | 0.54 | 0.03 | 0.02 | 0.08 | 0.05 | 0.33 | 0.50 |
| Melloul | 0.13 | 0.05 | 0.05 | 0.3 | - | **1.02** | 0.13 | 0.08 | 0.07 | 0.03 | 0.58 | 0.04 | 0.04 | 0.07 | 0.05 | 0.22 | 0.34 |
| Lakhdar | 0.18 | 0.05 | 0.06 | 0.9 | 0.70 | - | 0.39 | 0.29 | 0.21 | 0.16 | 0.80 | 0.20 | 0.18 | 0.30 | 0.15 | 0.66 | 2.30 |
| Tessaout | 0.01 | 0.01 | 0.01 | 0.08 | 0.17 | 0.34 | - | 0.08 | 0.03 | 0.01 | 0.33 | 0.02 | 0.02 | 0.06 | 0.04 | 0.23 | 0.17 |
| Isli | 0.02 | 0.01 | 0.02 | 0.06 | 0.17 | 0.24 | 0.05 | - | 0.10 | 0.06 | 0.41 | 0.08 | 0.07 | 0.35 | 0.08 | 0.99 | 0.27 |
| Ifni | 0.15 | 0.04 | 0.04 | 0.17 | **2.16** | 0.63 | 0.12 | 0.14 | - | 0.33 | **1.65** | 0.06 | 0.05 | 0.07 | 0.15 | 0.58 | 0.15 |
| Tifnoute | 0.005 | 0.01 | 0.01 | 0.02 | 0.73 | 0.26 | 0.03 | 0.03 | **1.69** | - | 0.79 | 0.00 | 0.00 | 0.03 | 0.09 | 0.51 | 0.12 |
| Tamda | 0.33 | 0.09 | 0.09 | 0.51 | **1.40** | **1.27** | 0.31 | 0.34 | **2.39** | **1.17** | - | 0.66 | 0.61 | 0.56 | 0.82 | **1.45** | 0.55 |
| Farda | 0.00 | 0.01 | 0.01 | 0.03 | **1.09** | 0.39 | 0.03 | 0.04 | **3.50** | - | **1.75** | - | - | 0.08 | 0.10 | 0.59 | 0.14 |
| Kannar | 0.00 | 0.01 | 0.01 | 0.03 | **1.01** | 0.36 | 0.03 | 0.04 | **2.96** | - | **1.62** | - | - | 0.07 | 0.09 | 0.09 | 0.13 |
| Rheraya | 0.07 | 0.03 | 0.04 | 0.14 | 0.45 | 0.52 | 0.10 | 0.13 | 0.60 | 0.17 | 0.94 | 0.27 | 0.25 | - | 0.093 | 0.61 | 0.66 |
| Ourika | 0.01 | 0.01 | 0.01 | 0.03 | 0.36 | 0.20 | 0.03 | 0.04 | 0.44 | 0.09 | 0.96 | 0.09 | 0.09 | 0.17 | - | 0.52 | 0.81 |
| Moulouya | 0.56 | 0.05 | 0.06 | 0.24 | 0.74 | 0.76 | 0.17 | 0.20 | **2.48** | 0.52 | **3.51** | 0.85 | 0.78 | 0.67 | 0.41 | - | 0.46 |
| Rachid | 0.00 | 0.003 | 0.01 | 0.01 | 0.16 | 0.14 | 0.01 | 0.02 | 0.13 | 0.004 | 0.25 | 0.00 | 0.00 | 0.05 | 0.01 | 0.28 |  |

**Table S9**. Heterozygosity excess and mode-shift distribution for Moroccan trout populations. IAM = infinitive allele model; SSM = stepwise mutation model; TPM = two-phase model. Bold values indicate a positive signal of bottleneck under a sign-rank Wilcoxon test.

| Population | Loci with heterozygosity excess | | | Mode-shift distribution |
| --- | --- | --- | --- | --- |
|  | **IAM** | **SSM** | **TPM** |  |
| Sidi Hamza | 0.320 | 0.578 | 0.843 | Normal L-shaped |
| Isli | 1.000 | 1.000 | 1.000 | Normal L-shaped |
| Melloul | 0.891 | 0.969 | 0.984 | Normal L-shaped |
| Miaami | 0.191 | 0.371 | 0.527 | Shifted |
| Lakhdar | 0.473 | 0.726 | 0.980 | Normal L-shaped |
| Farda | 0.326 | 0.633 | 0.752 | Normal L-shaped |
| Kannar | 0.937 | 1.000 | 1.000 | Normal L-shaped |
| Moulouya | **0.027** | 0.234 | 0.765 | Normal L-shaped |
| Tamda | 0.320 | 0.629 | 0.808 | Normal L-shaped |
| Ourika | 0.125 | 0.125 | 1.000 | Normal L-shaped |
| Rheraya | **0.008** | 0.500 | 0.500 | Shifted |
| Ifni | 0.422 | 0.781 | 0.922 | Normal L-shaped |
| Tifnoute | **0.008** | **0.008** | **0.008** | Normal L-shaped |
| Tessaout | 0.937 | 1.000 | 1.000 | Normal L-shaped |
| Dades | 0.812 | 0.812 | 0.875 | Normal L-shaped |
| M’Goun | 0.844 | 0.844 | 0.906 | Normal L-shaped |
| Sidi Rachid | **0.01** | 0.098 | 0.578 | Shifted |

**Table S10**. Number of trout released in particular Moroccan river systems over a 10-year period (2004–2014). Source: HCEFLCD, 2014^1^.

| POPULATION | 2004 | 2005 | 2006 | 2007 | 2008 | 2009 | 2010 | 2013 | 2014 |
| --- | --- | --- | --- | --- | --- | --- | --- | --- | --- |
| Sidi Rachid R. | 20,000 | 50,000 | 20,000 | 20,000 | 10,000 | 20,000 | 2,000 | 10,000 |  |
| Amengouss R. | 6,000 | - | - | - | 14,000 | - | - | - | - |
| Miaami R. | 10,000 | 10,000 | - | - | - | - | - | - | - |
| Oum er Rbia Basin | - | - | - | 20,000 | 10,000 | - | - | - | - |
| Talambote R. (Farda Basin) | - | - | - | 800 | - | - | 32,000 | - | - |

1. HCEFLCD. Rapports annuels de la pêche dans les eaux continentales, 2004-2014.

**Figure S1**. Mismatch distribution graphics for all analysed populations based on both mitochondrial markers (*MT-CYB* and D-loop).

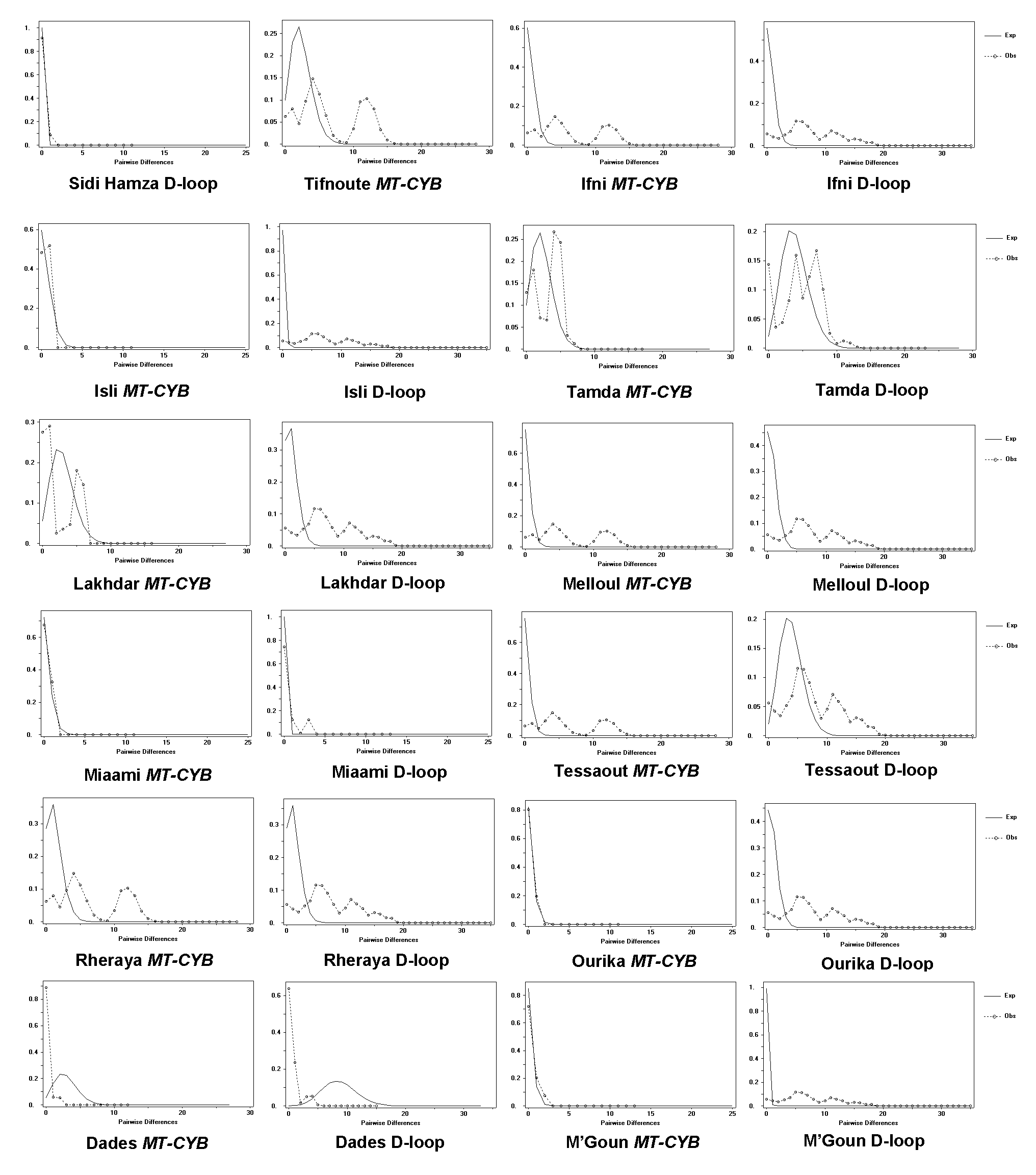
